# Optical nanosensor passivation enables highly sensitive detection of the inflammatory cytokine IL-6

**DOI:** 10.1101/2023.05.10.540217

**Authors:** Pooja Gaikwad, Nazifa Rahman, Rooshi Parikh, Jalen Crespo, Zachary Cohen, Ryan Williams

## Abstract

Interleukin-6 (IL-6) is known to a play critical role in the progression of inflammatory diseases such as cardiovascular disease, cancer, sepsis, viral infection, neurological disease, and autoimmune diseases. Emerging diagnostic and prognostic tools, such as optical nanosensors, experience challenges in successful clinical application in part due to protein corona formation dampening their selectivity and sensitivity. To address this problem, we explored the rational screening of several classes of biomolecules to be employed as agents in non-covalent surface passivation as a strategy to screen interference from non-specific proteins. Findings from this screening were applied to the detection of IL-6 by a fluorescent antibody-conjugated single-walled carbon nanotube (SWCNT)-based nanosensor. The IL-6 nanosensor exhibited highly sensitive and specific detection after passivation with a polymer, poly-L-lysine, as demonstrated by IL-6 detection in human serum within a clinically relevant range of 25 pg/mL to 25,000 pg/mL, exhibiting a limit of detection over three orders of magnitude lower than prior antibody-conjugated SWCNT sensors. This work holds the potential for rapid and highly sensitive detection of IL-6 in clinical settings with future application to other cytokines or disease-specific biomarkers.

## Introduction

Nanobiosensors have substantial potential in medical diagnostics^1^. Single-walled carbon nanotube (SWCNT)-based optical nanosensors have garnered interest due to the unique characteristics of near-infrared SWCNT photoluminescence, very little of which is absorbed by biological fluids and tissues^2-8^. Single-walled carbon nanotubes exhibit a range of chiralities described by (n, m) indices, each having discrete and narrow absorption and fluorescence emission bands, as well as large Stokes shifts^9, 10, 11^. Furthermore, the fluorescence of SWCNT does not decrease or photobleach over time due to excitation, allowing for long-term and frequent imaging and interrogation^12^.

The optoelectronic properties of SWCNT are affected by the surrounding environment, which makes them suitable for biosensing applications. Single-stranded (ss) DNA-wrapped SWCNTs have been used as optical nanosensors for cancers and metabolic diseases, among others^13-17^.

These optical nanosensors allow rapid and inexpensive detection compared to traditional techniques such as ELISA, mass spectrometry, and others^18, 19^. For example, simultaneous detection in human serum of multiple biomarkers for hepatocellular carcinoma, alpha fetoprotein and Golgi protein 73, has been reported^20^. Antibody-conjugated SWCNT-based optical sensors for cardiovascular disease biomarkers^19^, atherosclerosis^21^, cancer^5, 22, 23^, diabetes^24, 25^, and recently COVID-19^26^ have also been reported. Furthermore, successful in vivo detection of protein, lipid, nucleic acid, and small molecule analytes has been demonstrated, for example human epididymis protein 4, an FDA approved biomarker for high-grade serous ovarian carcinoma (HGSC)^4^, amongst others^27-31^.

Interleukin-6 (IL-6) is known to play a critical role in a wide range of diseases such as cardiovascular diseases^32-34^, viral infections such as COVID^35^, cancer^36, 37^, sepsis and bacterial infection^38^, neurological diseases^38^, and autoimmune diseases^39, 40^. Therapies which reduce the effects of IL6 have significantly improved treatment outcomes for rheumatoid arthritis and COVID-19^35, 41^. The timely determination of elevated IL-6 levels offers clinicians an opportunity to identify the individuals at risk for poor outcomes and treat them appropriately. Standard techniques for detection of IL-6 are immunoassays, western blotting, mass spectroscopy, flow cytometry^42^, and semi-quantitative immunohistochemical detection^43^. Of these, immunosorbent assays offer the lowest limits of IL-6 detection in human plasma, as low as 0.5 pg/mL^44, 45^.

However, these techniques require long incubation periods, trained personnel, and proper instrumentation, making them incompatible for routine clinical diagnosis, frequent testing, and monitoring^46, 47^. To mitigate these limitations, immunosensing antibody-based nanosensors have emerged as a user-friendly point-of-care alternative^46^. For example, nanosensors for IL-6 based on silica nanowires and metal nanoparticles have been reported^48-51^.

To successfully implement the clinical use of SWCNT-based nanosensors, their sensitivity and selectivity must be retained in complex biological fluids. The formation of a heterogenous protein corona on the nanotube surface in physiologically relevant environments limits this potential^27, 52-54^. This phenomenon is due to the hydrophobicity nanotubes, leading to adsorption of proteins and other biologics in a non-covalent and non-specific manner^55, 56^. This leads to a reorientation of the dipole moments and charge transfer around ssDNA-SWCNTs, leading to photoluminescence modulation^57^. Therefore, in order to prevent non-specific nanosensor responses to the detriment of selectivity and sensitivity, it is necessary to prevent such interactions^14^.

As non-specific interactions with serum components are entropically favored, these coronas are difficult to remove once formed. However, it is possible to saturate the nanotube surface with a homogenous, known corona prior to nanotube interaction with serum components. Non-covalent surface adsorption prior to sensor deployment, here referred to as passivation, by biocompatible molecules is a widely used strategy for controlling the surface coronas of materials, including nanosensors and standard molecular biology assays such as immunohistochemistry, Western blotting, and immunoassays^4, 5, 58-62^. Such strategies are particularly prevalent in assays involving the use of antibodies as molecular recognition elements.

Serum albumin and polyethylene glycol (PEG)-modified phospholipids were used in previous studies as passivation agents to improve the performance of SWCNT-based optical nanosensors^4, 5, 14, 63^. Proteins such as bovine serum albumin (BSA) and non-fat dry milk (NFDM) have been used extensively in biological assays for passivation due to their low cost and biocompatibility^64, 65^. PEG-modified phospholipids exhibit unique biomimetic properties and have been used particularly for in vivo applications^66-68^. Polymers such as polyethylene imine^69^ (PEI), chlorin e6^70^, and poly-L-lysine^71^ (PLK) have been used to non-covalently adsorb to nanotube surfaces for imaging, rather than sensing, applications.

Here, we studied the efficacy of passivation agents in improving the performance of an IL-6 specific photoluminescent nanosensor in human serum. We screened 8 passivation agents across the general classes of proteins, polymers, and PEG-modified phospholipids for their potential to block non-specific adsorption of serum components onto the nanotube surface. Promising passivation molecules were then used in the improvement of an antibody-conjugated IL-6 nanotube-based optical sensor in human serum. We found that BSA and PLK, which has not previously been explored for nanosensors, impart pg/mL range detection of IL-6 in this complex biological environment.

### Experimental Section

#### Preparation of SWCNT-ssDNA

We initially sought to evaluate the potential of passivation agents using the basic nanotube construct, without a molecular recognition moiety, in order to simplify the screening system. HiPCO-prepared single-walled carbon nanotubes (SWCNT) (NanoIntegris Technologies, Inc.; Quebec, Canada) and a single-stranded DNA of the sequence (TAT)_6_ (Integrated DNA Technologies; Iowa, USA) in a 1:2 mass ratio were suspended in 0.5 mL 1x phosphate buffered saline (Sigma-Aldrich; Missouri, USA). The suspension was sonicated for 1 hour at 40% amplitude while on ice via a 120 W ultrasonicator with 1/8″ probe microtip (Fisher Scientific; New Hampshire, USA). The sonicated suspension was ultracentrifuged for 1 hour at 58,000 x g in 4 ml polycarbonate centrifuge tubes (Beckman Coulter; California, USA) using an Optima Max-XP Ultracentrifuge (Beckman Coulter) fit with an MLA-50 rotor. After ultracentrifugation, the top 75% of the suspension was collected and filtered on the day of use with 100 kDa Amicon Ultra 0.5 ml centrifugal filters (Sigma-Aldrich) at 14,000 x g for 15 minutes to remove unwrapped ssDNA. After filtration, the solution retained in the filter containing ssDNA-SWCNT was washed two times with 200 μL 1X phosphate buffer saline (PBS) and centrifugal-filtered again. Finally, the solution containing SWCNT-(TAT)_6_ retained in the filter was suspended in total 200 uL of 1x PBS.

The concentration of SWCNT-(TAT)_6_ was determined by using a V-730 UV–visible absorption Spectrophotometer (Jasco Inc.; Maryland, USA). SWCNT-(TAT)_6_ solution was diluted with 1x PBS to obtain absorbance in the range of 0.3 to 0.7. The concentration of SWCNT was calculated using the value of the absorbance minima near 630 nm (Extinction coefficient = 0.036 L mg-1 cm-1)4, 5, 19.

#### Fluorescence spectroscopy for screening

Near-infrared fluorescence emission spectra of SWCNT-(TAT)_6_ were acquired in a MiniTracer spectrophotometer (Applied NanoFluorescence; Texas, USA). The laser source has an excitation wavelength of 638 nm and emission spectra were obtained between 900 nm and 1400 nm. SWCNT (7,5), (7,6), and (9,5) chiralities were analyzed using custom MATLAB code (available upon request) which fit emission peaks to a baseline-subtracted pseudo-Voigt model^4, 5, 72^. Fits were ensured to have goodness of fit values (R^2^) greater than 0.98 prior to analyses. Center wavelengths were obtained from each fit, as well as maximum intensity values.

#### Screening of passivation agents in complex biological media

Passivation of SWCNT-(TAT)_6_ complexes was achieved by incubating 0.5 mg/L SWCNT-(TAT)_6_ and passivating agents in the desired mass ratio at 4 °C for 30 minutes. Passivation agents used, including BSA, non-fat dry milk (NFDM), casein, PEG-1500, PEI, PLK, 16:0 PE PEG, and DSPE-PEG-amine (**Table 1**; Supplementary Figure S1), were evaluated in mass ratios of 5x, 25x, 50x and 100x greater than the nanotube complex. The molar ratio of protein passivation agents to SWCNT-(TAT)_6_ at a 50x mass ratio was 425:1 for BSA and 1409-1127:1 for casein; the molar ratio of polymer passivation agents to SWCNT-(TAT)_6_ at 50x mass ratio is 188-403:1 for PLK, 2819:1 for PEI and 12,000:1 for PEG; and the molar ratio of phospholipid passivation agents to SWCNT-(TAT)_6_ at a 50x mass ratio are 14,092:1 for DSPE-PEG (NH2) and 10,251:1 for 16:0 PE2000PEG (Supplementary Calculations). 10% heat inactivated fetal bovine serum (FBS) (Corning; New York, USA) was used to challenge each passivation, simulating complex in vivo biological conditions. Changes resulting from the addition of FBS to passivated SWCNT-(TAT)_6_ were quantified by a change in the fluorescence emission center wavelength for the (7,5), (7,6), and (9,5) chiralities. Data were collected at 2, 15, 30, 60, 120, 150, and 180 minutes after challenge with FBS and acquired in triplicate.

**Table 1.**
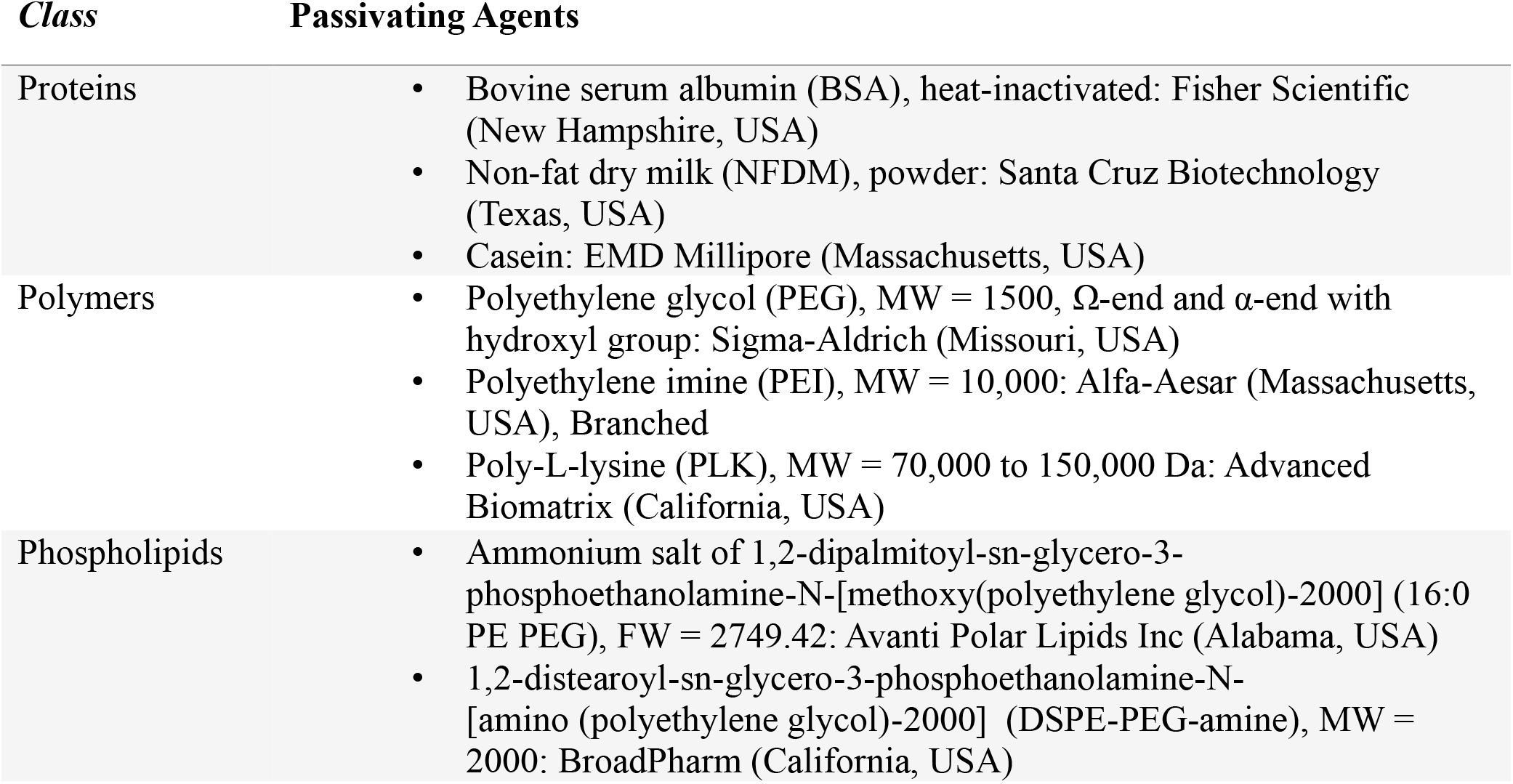
Passivation agents used in screening.

#### Absorption spectroscopy

Near-IR absorption spectra in the range of 900 nm to 1400 nm were acquired using a MiniTracer spectrophotometer to analyze the formation of stable interactions between passivation agents and SWCNT constructs. Data were analyzed using custom MATLAB code as above. Mass ratios of 5x, 10x, 25x, and 50x of the passivation agent in comparison to 10 mg/L SWCNT-(TAT)_6_ were used. Changes in absorbance resulting from nanotube passivation were quantified by changes in the absorbance center wavelength peaks of nanotube chiralities (7,5), (7,6), (9,5), and (6,5). Data were collected at 2, 15, 30, 60, 120, 150, and 180 minutes post-passivation to confirm the duration for which passivation is stable. Data were acquired in triplicate.

#### IL-6 sensor synthesis

SWCNT-(TAT)_6_ was prepared as above with the modification that the ssDNA sequence contained a 3’ primary amine modification (5’-(TAT)_6_)/3AmMO/-3’) (Integrated DNA Technologies). The amine-modified ssDNA-SWCNT complex was conjugated to a monoclonal IL-6 antibody (Catalog number 554543; BD Biosciences, California, USA) using carbodiimide conjugation chemistry similarly to previous studies^4, 5^. Briefly, the carboxylic acid group of the antibodies were activated using 1-ethyl-3-(3-dimethylainopropyl)carbodiimide) (Sigma-Aldrich) and N-hydroxysuccinimide (TCI Chemicals, Oregon, USA), in a 10x and 25x molar excess respectively, for 15 min at 4 °C. The reaction was quenched with 1 uL of 2-mercaptoethanol (Sigma-Aldrich). The activated antibodies were mixed with SWCNT-(TAT)_6_-NH2 in 1:1 molar ratio of ssDNA to antibody. The reaction mixture was incubated at 4 °C on ice for a total of 2 hours, with gentle brief vortexing every 30 minutes. The reaction mixture was dialyzed against deionized water with a 1,000 kDa MWCO filter (Float-A-Lyzer G2; Spectrum Labs, California, USA) at 4 °C for 48 hours with two dialysate changes.

#### Physicochemical characterization of the IL-6 nanosensor

To confirm successful conjugation of antibody to the ssDNA-nanotube construct, we performed light scattering measurements. Dynamic light scattering (DLS) was performed (Nano-ZS90, Malvern: Worcestershire, United Kingdom) for nanosensor and SWCNT-(TAT)_6_-NH2 nanotube complexes as previously described to determine their relative sizes^4, 5^. Electrophoretic light scattering (ELS) (Nano-ZS90, Malvern) was performed to compare the relative ζ -potential of the nanosensor and SWCNT-(TAT)_6_-NH2 complex. Data were acquired in triplicate.

#### Evaluation of IL-6 nanosensor function

To confirm basic functionality of the IL-6 nanosensor complex, we first evaluated the fluorescence response of 0.5 mg/L nanosensor to 5250 ng/mL human IL-6 (Catalog number RP8619; Thermo Fisher Scientific, Massachusetts, USA) in 1x PBS using the MiniTracer as described above with identical timepoints and data processing. The response of the sensor passivated with 50x BSA as in prior studies was analyzed in 10% Fetal Bovine Serum as described above. All experiments were performed in triplicate.

#### IL-6 nanosensor in human serum

0.5 mg/L IL-6 nanosensor was passivated with 50x molar mass ratios of PLK, NFDM, and BSA at 4 °C for 30 minutes. Fluorescence data were acquired before and after adding pooled human serum (MP Biomedicals; California, USA) with or without spiked IL-6 using a custom-built high-throughput near-IR plate reader spectrophotometer (ClaIR, Photon Etc.; Montreal, Canada). Spiked IL-6 protein was added to final concentrations of 25 pg/mL, 250 pg/mL, 2,500 pg/mL, and 25,000 pg/mL. Control responses were obtained with the nanosensor construct without passivation. Data were acquired in triplicate. Custom MATLAB code was used to analyze and fit individual nanotube chirality peaks to a pseudo-Voigt model (code available upon request). The analyzed fluorescence emission chiralities were (7,5), (7,6), and (9,5) for excitation with the 655 nm laser source and (10,2), (9,4), (8,6), (8,7) for excitation with the 730 nm laser source.

#### Statistical analysis and data processing

All statistical analyses were performed in OriginPro 2021 (OriginLab Corporation; Massachusetts, USA). Passivation agent screening experiments were analyzed by one way analysis of variance (ANOVA) with Dunnett post-hoc analysis. P-values were assigned **** = P < 0.0001, *** = P < 0.001, ** = P < 0.01, and * = P < 0.05. Zeta potential results for IL-6 nanosensor characterization and in vitro performance of IL-6 nanosensor were analyzed by a two-tailed t-test. Experiments in human serum were analyzed via one way ANOVA with Dunnet post-hoc analysis to compare to the non-passivated control.

Changes in SWCNT center wavelength were reported relative to their emission prior to analyte addition. Custom MATLAB code used for data processing and analysis is available upon request.

## Results and Discussion

### Interference caused by non-specific proteins to nanosensors

Fetal bovine serum is commonly used in sensor development studies as a model complex protein environment^73^. We assessed the non-specific interactions of FBS with SWCNT-(TAT)_6_ by evaluating shifts in the center wavelength of the E11 peaks in absorbance and fluorescence spectra. Fluorescence peaks exhibited substantial shifts in the presence of 10% FBS. The shifts in the (7,5), (7,6), and (9,5) chiralities were, respectively, red-shifted (bathochromic) 2.3 ± 0.37 nm, 7.2 ± 0.24 nm, and blue shifted (hypsochromic) 2.5 ± 0.7 nm (**Figure 1A**). Absorbance peaks exhibited center wavelength shifts of 3.22 ± 0.6 nm, 2.3 ± 0.23 nm, 3.6 ± 0.2 nm, and 3.7 ± 0.08 nm for (6,5), (7,5), (7,6), and (9,5) nanotube chiralities respectively (**Figure 1B**). We hypothesized that passivation agents which can reduce the magnitude of photoluminescence changes upon challenging with 10% FBS would improve the sensitivity and specificity of physiologically relevant nanosensors.

**Figure 1.**
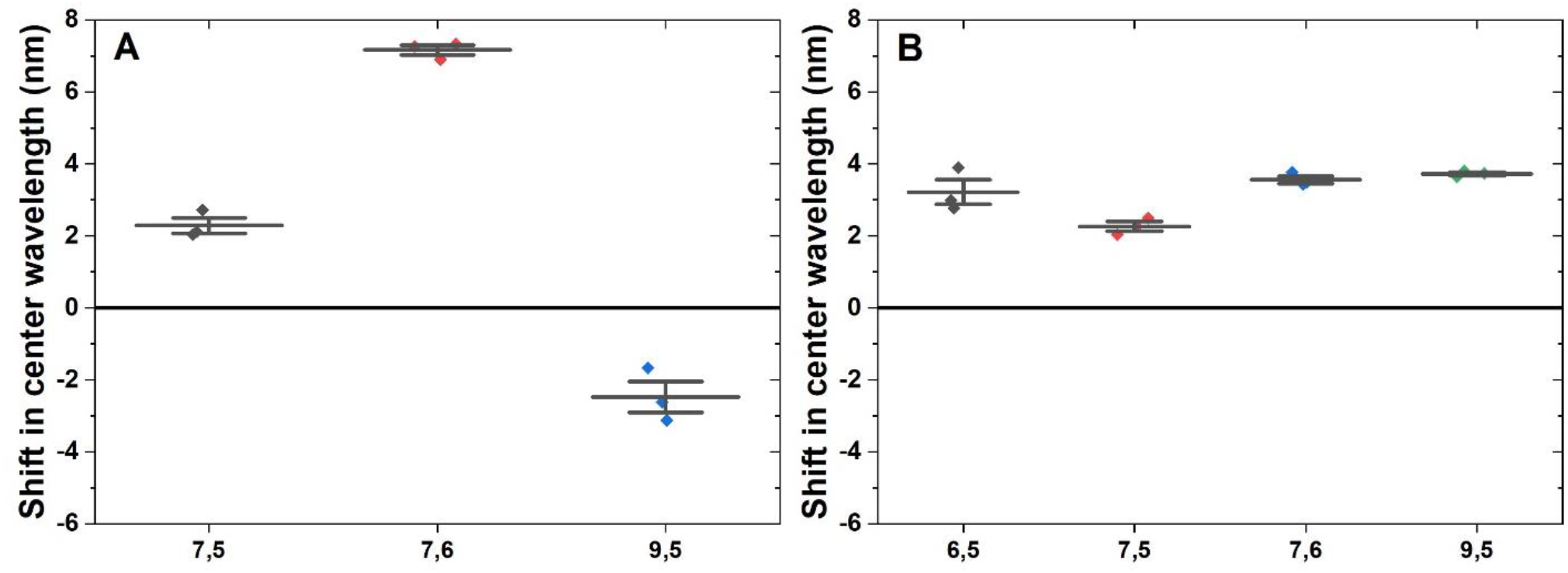
Non-specific interactions of 10% FBS with SWCNT-TAT_6_. (A) Shift in center wavelength of the E11 fluorescence peaks for chiralities (7,5) (2.3 ± 0.37 nm), (7,6) (7.2 ± 0.2 nm), and (9,5) (−2.5 ± 0.7 nm) (B) Shift in center wavelength of the E11 absorbance peak for chiralities (6,5) (3.2 ± 0.6 nm), (7,5) (2.3 ± 0.2 nm), (7,6) (3.6 ± 0.2 nm), and (9,5) (3.7 ± 8E-2). N= 3.

### Proteins as passivating agents

Previous studies have shown that BSA successfully screened interference caused by non-specific proteins and improved selectivity of the nanosensors^4, 5^. Here, we explored higher and lower mass ratios compared to the previously-used 50x mass ratio. Three hours after challenging with FBS, we observed a shift of 7.9 ± 2.2 nm for 5x, 9.6 ± 0.5 nm for 25x, 4.6 ± 0.26 nm for 50x, and 7.7 ± 3.3 nm for 100x for the (7,6) fluorescence peak (**Figure 2A**). Across all ratios, upon challenging with FBS, BSA passivation did not significantly mitigate FBS-induced shifting. Similar observations were made for the (7,5) and (9,5) fluorescence peaks (Supplementary Figures S2, S3).

**Figure 2.**
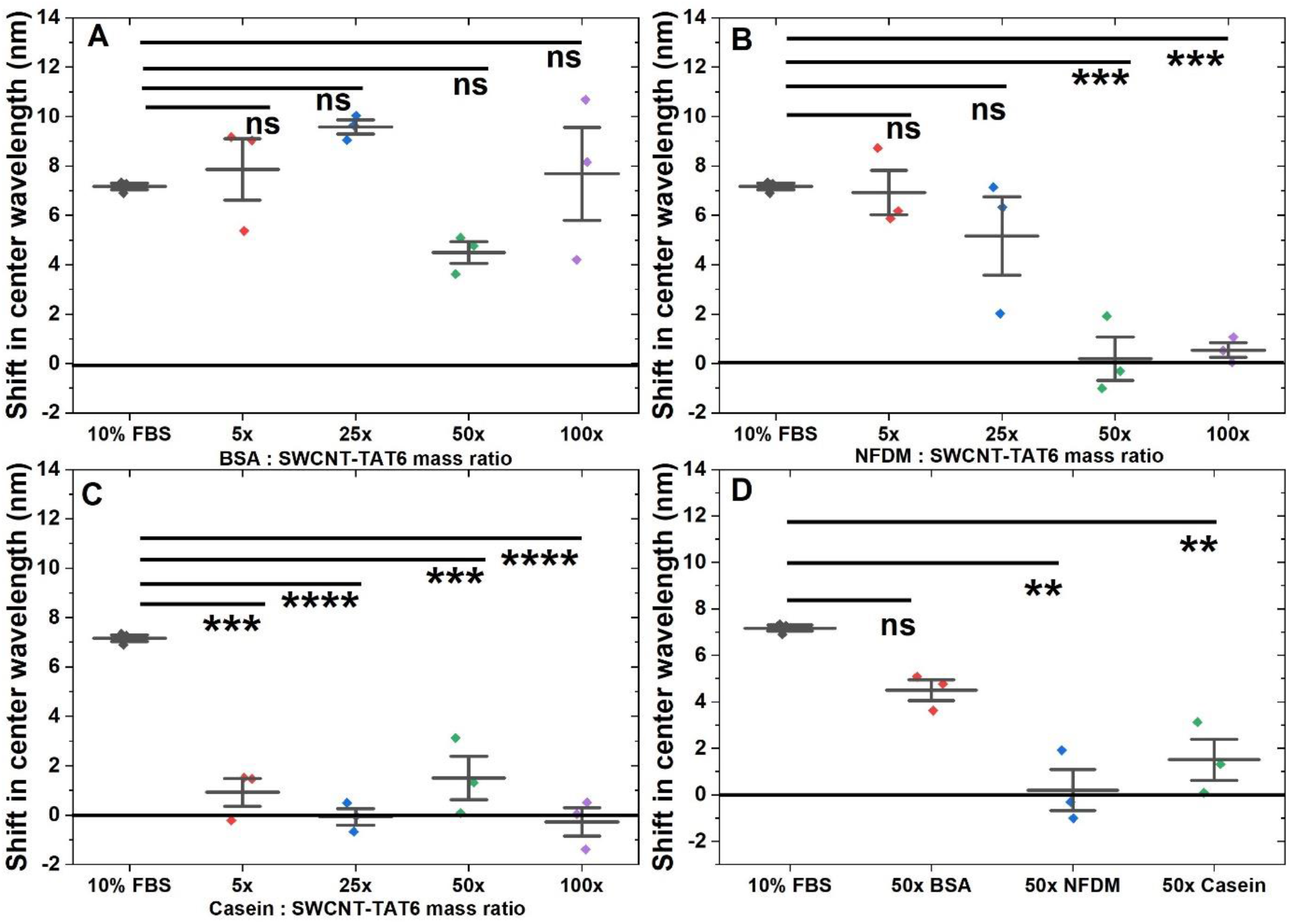
Screening serum interference with protein passivation of SWCNT-(TAT)_6_. Shift in emission center wavelength for (7,6) upon adding FBS (A) for BSA passivation, n=3, mean ± SD. 10% FBS (7.2 ± 0.2 nm), 5x BSA (7.9 ± 2.2 nm), 25x BSA (9.6 ± 0.5 nm), 50x BSA (4.6 ± 0.3 nm), 100x BSA (7.7 ± 3.3 nm). The difference in means for groups: FBS and 5x BSA (0.69 nm, p=0.94), FBS and 25x BSA (2.4 nm, p=0.25), FBS and 50x BSA (2.7 nm, p=0.18), FBS and 100x BSA (0.5 nm, p=0.98). (B) for NFDM passivation, n=3, mean ± SD. 10% FBS (7.2 ± 0.2 nm), 5x NFDM (7 ± 1.6 nm), 25x NFDM (5.2 ± 2.7 nm), 50x NFDM (0.2 ± 1.5 nm), 100x NFDM (0.5 ± 0.5nm). Difference in means for groups: FBS and 5x NFDM (0.24 nm, p=0.998), FBS and 25x NFDM (2 nm, p=0.24), FBS and 50x NFDM (6.97 nm, p=5.2E-4), FBS and 100x NFDM (6.6 nm, p=7.3E-4) (C) for Casein passivation, n=3, mean ± SD. 10% FBS (7.2 ± 0.2 nm), 5x casein (0.9 ± 1 nm), 25x casein (−0.1 ± 0.6 nm), 50x casein (1.5 ± 1.5 nm), 100x casein (−0.3 ± 1 nm). Difference in means for groups: FBS and 5x Casein (6.2 nm, p=1.85E-4), FBS and 25x Casein (7.24 nm, p=7.1E-5), FBS and 50x Casein (5.7 nm, p=3.8E-4), FBS and 100x Casein (7.45 nm, p=4.91E-5) (D) Comparison of passivation agents at the 50x mass ratio, shift in emission center wavelength n=3, mean ± SD: FBS and 50x BSA (2.7 nm, p=0.1); FBS and 50x NFDM (7 nm, p=1.5E-3); FBS and 50x Casein (5.7 nm, p=4.5E-3).

We further explored other protein passivation agents, including non-fat dry milk (NFDM) and its primary component, casein protein. We found that NFDM passivation mitigated shifting at the 50x mass ratio (**Figure 2B**). We observed that a 50x mass ratio was optimal for NFDM and casein passivation, with shifts of 0.2 ± 1.5 nm (**Figure 2B**) and 1.5 ± 1.5 nm (**Figure 2C**), respectively, for the (7,6) chirality. A 50x mass ratio was optimal for interference screening of the (7,5) and (9,5) chiralities as well (Supplementary Figures S2, S3). For casein passivation, as low as 5x mass ratio was found to be effective for all three chiralities investigated (**Figure 2C**; Supplementary Figures S2, S3), whereas NFDM passivation was not effective at this ratio (**Figure 2B**; Supplementary Figures S2, S3). As casein is just one component of NFDM^74^, this may contribute to the difference in effectiveness at various mass ratios. The response for all three observed chiralities remained relatively stable for 180 minutes in the presence of each passivation agent (Supplementary Figures S4, S5, S6).

### Polymers as passivation agents

We next investigated several polymers as potential passivating agents. We evaluated two cationic polymers, poly-L-lysine (PLK) and polyethyleneimine (PEI), and one anionic polymer, 1500 Da polyethylene glycol (PEG). PEI and PLK have been used to noncovalently wrap SWCNTs for various applications, though not as passivating agents^69, 71^. PEG has been used to covalently functionalize SWCNTs and was chosen to help us distinguish the effect of charge-based interactions on ssDNA-wrapped SWCNTs. We hypothesized that cationic polymers would be better candidates for passivation due to attractive ionic interactions with anionic ssDNA.

PLK demonstrated a decrease in FBS interference across all passivation ratios investigated (**Figure 3A**). Branched 10 kDa polyethylene imine (PEI) demonstrated a decrease in interference caused by FBS for all ratios investigated (**Figure 3B**), though significant visible flocculation of the nanotube construct was observed (Supplementary Figure S7). It is likely that this results from the branched nature of the PEI used, with multiple amine groups, bringing multiple nanotube constructs physically closer together. Passivation of the nanotube construct with PLK resulted in minimal shifts and no flocculation at all mass ratios investigated upon challenging with FBS, with all shift magnitudes ≤0.3 nm (**Figure 3A**; Supplementary Figures S8, S9). We then evaluated PEG-1500, an anionic polymer, demonstrating no screening effect against FBS at all ratios (**Figure 3C**; Supplementary Figures S8, S9). It is likely that the repulsive columbic interactions between anionic ssDNA and anionic PEG results in poor interaction with the nanotube surface, which in turn allows FBS components to interact with SWCNT-(TAT)_6_ non-specifically. PLK passivation showed no significant screening effect for the (9,5) chirality due to high variability, though it did for the (7,6) and (7,5) chiralities, indicating that a given passivation agent may show effective passivation at some, but not all, chiralities (**Figure 3A**; Supplementary Figures S8, S9). The response for all three fluorescence peaks for all three passivation agents remained relatively stable for a period of 180 minutes following addition of FBS as a challenger (Supplementary Figures S10, S11, S12).

**Figure 3.**
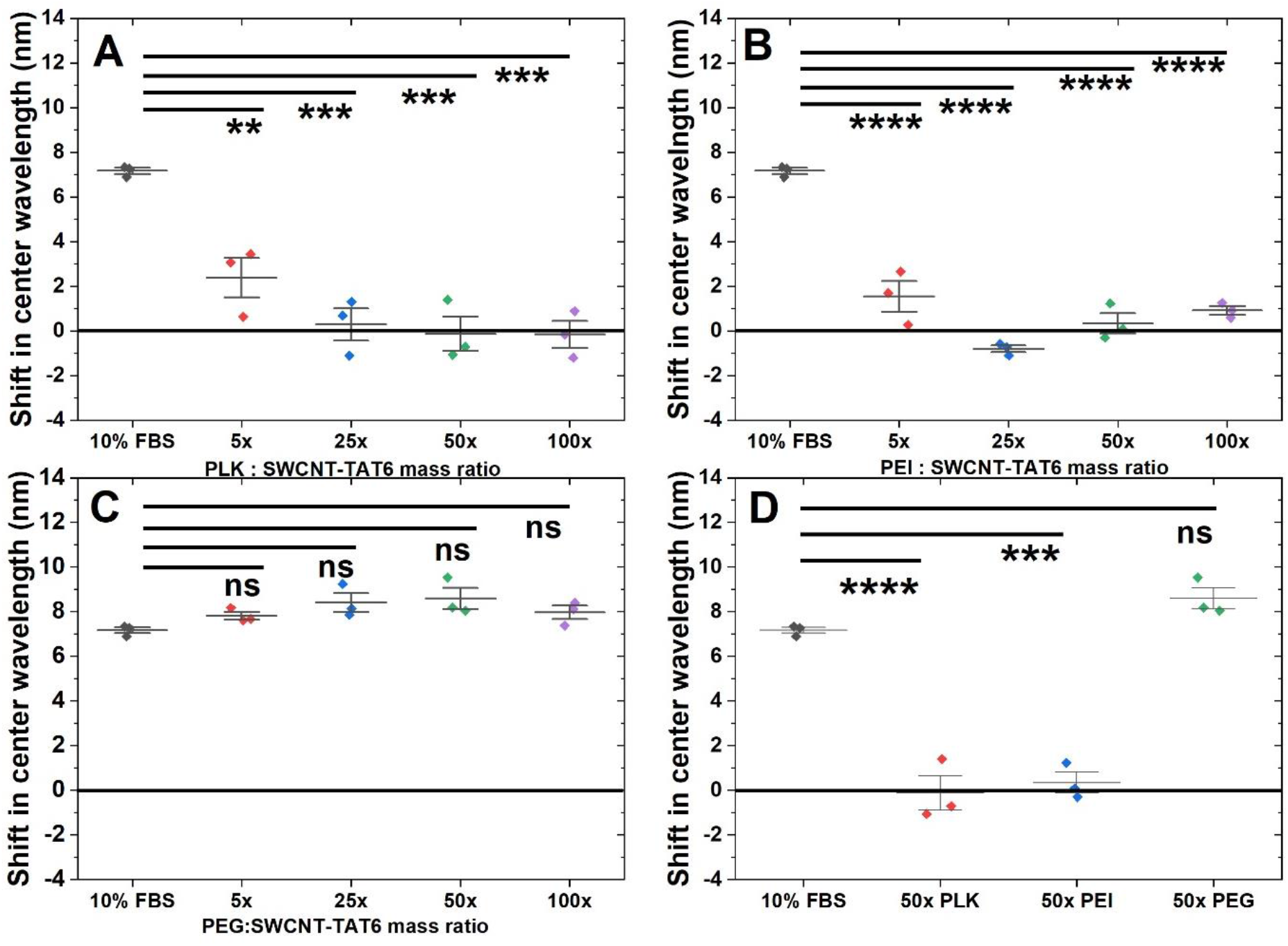
Screening serum interference with polymer passivation of SWCNT-(TAT)_6_. Shift in emission center wavelength for (7,6) upon adding FBS (A) for PLK passivation, n=3, mean ± SD. 10% FBS (7.2 ± 0.2 nm), 5x PLK (2.4 ± 1.5 nm), 25x PLK (0.3 ± 1.3 nm), 50x PLK (−0.13 ± 1.3 nm), 100x PLK (−0.2 ± 1 nm). Difference in means for groups: FBS and 5x PLK (4.8 nm, p=5.2E-3), FBS and 25x PLK (6.9 nm, p=5.14E-4), FBS and 50x PLK (7.3 nm, p=3.41E-4), FBS and 100x PLK (7.3 nm, p=3.31E-4) (B) for PEI passivation, n=3, mean ± SD. 10% FBS (7.2 ± 0.2 nm), 5x PEI (1.5 ± 1.2 nm), 25x PEI (−0.8 ± 0.3 nm), 50x PEI (0.3 ± 0.8 nm), 100x PEI (0.9 ± 0.3 nm). Difference in means for groups: FBS and 5x PEI (5.6 nm, p=2.13E-5), FBS and 25x PEI (8 nm, p=1.61E-6), FBS and 50x PEI (6.8 nm, p=7.43E-6), FBS and 100x PEI (6.2 nm, p=1.4E-5) (C) for PEG passivation, n=3, mean ± SD. 10% FBS (7.2 ± 0.2 nm), 5x PEG (7.8 ± 0.3 nm), 25x PEG (8.4 ± 0.7 nm), 50x PEG (8.6 ± 0.8 nm), 100x PEG (8 ± 0.5 nm). Difference of means for groups: FBS and 5x PEG (0.64 nm, p=0.49), FBS and 25x PEG (1.2 nm, p=0.09), FBS and 50x PEG (1.4 nm, p=0.053), FBS and 100x PEG (0.8 nm, p=0.35) (D) Comparison of passivation agents at a 50x mass ratio, shift in emission center wavelength n=3, mean ± SD. FBS and 50x PLK (7.3 nm, p=6.6E-5); FBS and 50x PEI (6.8 nm, p=1.5E-4); FBS and 50x PEG (1.4 nm, p=0.24).

### Phospholipids as passivation agents

We next evaluated two PEGylated phospholipids, 16:0 PE2000PEG and DSPE PEG (NH2). 16:0 PE2000PEG has been used to successfully passivate SWCNT-based nanosensors and to prepare SWCNT suspensions^63, 68^. All ratios of DSPE PEG (NH2) passivation showed no significant change in center wavelength compared to controls for all three chiralities investigated (**Figure 4A**; Supplementary Figures S13, S14). For 16:0 PE2000PEG passivation, 50x and 100x mass ratio passivation showed screening of FBS interference for the (7,6) and (9,5) chiralities only (**Figure 4B**; Supplementary Figures S13, S14). Again, the responses for most chiralities remained relatively stable for up to 180 minutes (Supplementary Figures S15, S16). For agents indicating successful screening of FBS interference, the mass ratios 50x and 100x gave comparable passivation results. Therefore, we concluded that a mass ratio of 50x is optimal for those agents as to not saturate the system with unnecessary components.

**Figure 4.**
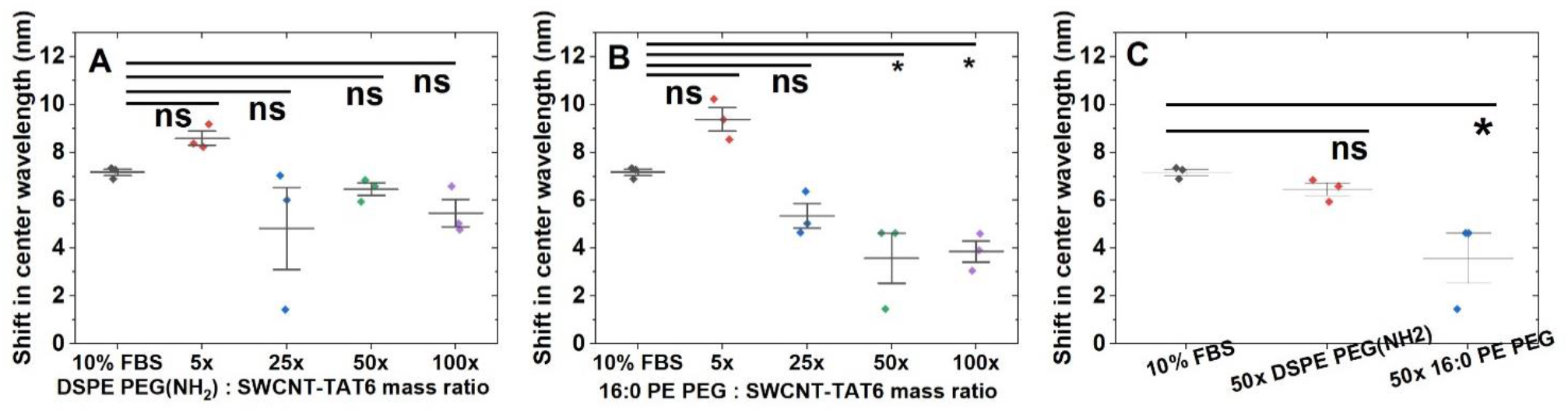
Screening serum interference with phospholipid passivation of SWCNT-(TAT)_6_. Change in emission center wavelength upon challenging with FBS. Shift in emission center wavelength for (7,6) upon adding FBS (A) for DSPE PEG (NH2) passivation, n=3, mean ± SD. 10% FBS (7.2 ± 0.2 nm), 5x DSPE PEG (NH2) (8.6 ± 0.5 nm), 25x DSPE PEG (NH2) (4.8 ± 3 nm), 50x DSPE PEG (NH2) (6.5 ± 0.5 nm), 100x DSPE PEG (NH2) (5.5 ± 1 nm). Difference in means for groups: FBS and 5x DSPE PEG (NH2) (1.4 nm, p=0.62), FBS and 25x DSPE PEG (NH2) (2.34, p=0.25), FBS and 50x DSPE PEG (NH2) (0.70 nm, p=0.94), FBS and 100x DSPE PEG (NH2) (1.71 nm, p=0.48). (B) for 16:0 PE PEG passivation, n=3, mean ± SD. 10% FBS (7.2 ± 0.2 nm), 5x 16:0 PE PEG (9.4 ± 0.8 nm), 25x 16:0 PE PEG (5.4 ± 0.9 nm), 50x 16:0 PE PEG (3.6 ± 1.8 nm), 100x 16:0 PE PEG (3.9 ± 0.8 nm). Difference in means for groups: FBS and 5x 16:0 PE PEG (2.21 nm, p=0.13), FBS and 25x 16:0 PE PEG (1.81 nm, p=0.23), FBS and 50x 16:0 PE PEG (3.6 nm, p=1.5E-2), FBS and 100x 16:0 PE PEG (3.3 nm, p=2.3E-2) (C) Comparison of passivation agents at a 50x mass ratio, shift in emission center wavelength n=3, mean ± SD. FBS and 50x DSPE PEG (NH2) (0.7, p=0.7); FBS and 50x 16:0 PE PEG (3.6 nm, p=3.6E-2).

### Analysis of corona formation via absorbance spectroscopy

Changes in the proximal environment of SWCNT-(TAT)_6_ due to close interactions and/or homogenous corona formation with the passivating agents may be reflected by changes in the E11 absorbance maxima of each chirality^75-77^. We hypothesized that the ability of the passivation agent to screen non-specific interactions is correlated to its ability to form a stable corona around the nanotube complex. We assumed that stronger interactions of a given passivation agent with the nanotube surface would result in greater changes in the center wavelength of SWCNT absorption peaks. Further, we propose that to successfully screen non-specific interactions by 10% FBS, the passivation agent should interact with SWCNT-(TAT)_6_ more strongly than FBS itself. Therefore, we anticipated that passivation agents which result in larger absorbance shifts than that caused by 10% FBS would exhibit strong passivation capability. For this reason, changes in individual chirality absorption center wavelengths in the presence of 10% FBS were compared to those in the presence of passivation agents.

For non-passivated SWCNT-(TAT)_6_ in serum conditions, the center wavelength of the (7,6) chirality was shifted by 3.6 ± 0.2 nm as compared to buffer conditions (**Figure 1B**). We found that, at 180 minutes post-passivation with 50x BSA, the (7,6) emission peak center wavelength shifted by 1.32 ± 0.4 nm, smaller than 3.6 ± 0.2 nm for 10% FBS (**Figure 5A**). We observed similar responses for the (7,5), (9,5), and (6,5) chiralities (Supplementary Figures S17A, S18A, S19A). A significant shift was observed after addition of 50x NFDM for the (7,6) and (9,5) chiralities of 4.6 ± 0.4 nm and 4.56 ± 0.35 nm, respectively (**Figure 5A**; Supplementary Figure S18A). Further, 50x casein passivation showed a higher shift magnitude of 5.7 ± 0.1 nm than for FBS alone (**Figure 5A**).

**Figure 5.**
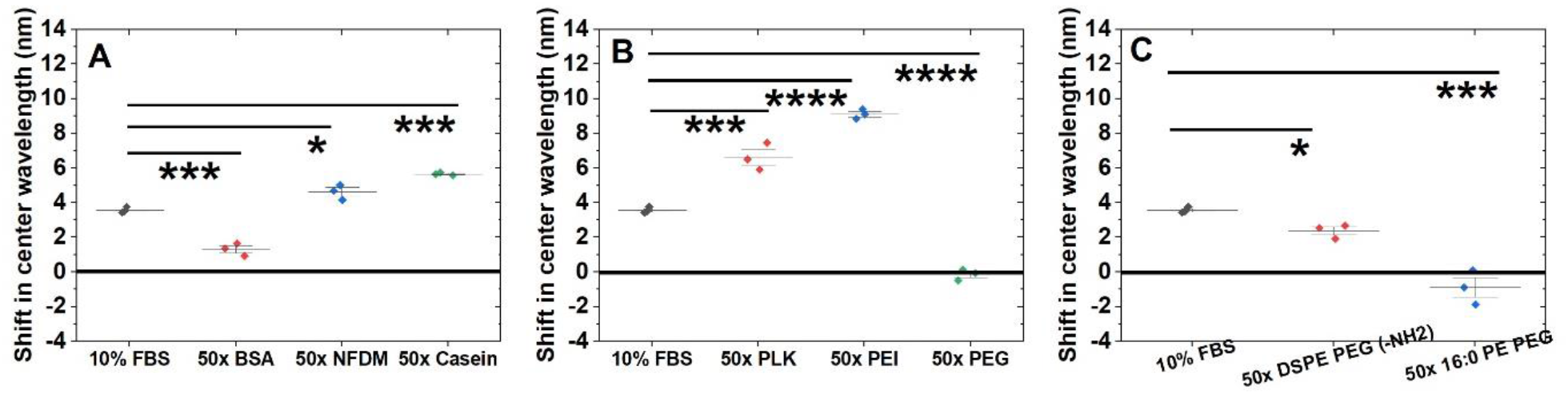
Change in absorbance as a measurement of homogenous and stable passivation. Comparison of the ability and to influence the proximal environment of SWCNT-(TAT)_6_ for (7,6) peak by (A) 50x mass ratio for protein passivation. n=3, mean ± SD. 10% FBS (3.6 ± 0.2 nm), 50x BSA (1.3 ± 0.4 nm), 50x NFDM (4.6 ± 0.4 nm), 50x casein (5.7 ± 0.1 nm). Comparison in means for groups: FBS and 50x BSA (2.2 nm, 4.1E-4), FBS and 50x NFDM (1.1 nm, p=1.85E-2); FBS and 50x Casein (2.1 nm, p=6.2E-4) (B) 50x mass ratio polymers. n=3, mean ± SD. 50x PLK (6.6 ± 0.8 nm), 50x PEI (9.6 ± 1 nm), 50x PEG (−0.1 ± 0.3 nm). Difference in means for groups: FBS and 50x PLK (3.1 nm, p=1.3E-4), FBS and 50x PEI (5.6 nm, p=5.2E-8), FBS and 50x PEG (3.69 nm, p=2.2E-5). (C) 50x mass ratio phospholipids, n=3, mean ± SD. 50x DSPE PEG (NH2) (2.38 ± 0.4 nm) and 50x 16:0 PE PEG (−0.9 ± 1 nm). Difference in means for groups: FBS and 50x DSPE PEG (NH2) (1.2 nm, p=0.08), FBS and 50x 16:0 PE PEG (4.4 nm, p=8.2E-4).

Of the polymeric passivation agents investigated, 50x PLK and 50x PEI demonstrated absorbance shifts of 6.6 ± 0.8 nm and 9.6 ± 1 nm, respectively (**Figure 5B**). A similar trend was observed for these two agents when evaluating the (7,5) and (9,5) chiralities (Supplementary Figures S17B, S18B). For 50x PEG passivation, the (7,6) absorption center wavelength shifted by –0.14 ± 0.3 nm (**Figure 5B**). For all other peaks, 50x PEG demonstrated a consistent blue shift compared to that of serum conditions (Supplementary Figures S17B, S18B, S19B).

Upon evaluation of absorption shifts induced by phospholipid agents, 50x DSPE PEG (NH2) caused a shift of 2.38 ± 0.4 nm, smaller than that of FBS (**Figure 5C**). For this chirality, 50x 16:0 PE PEG passivation caused a blue shift of 0.9 ± 1 nm (**Figure 5C**). Other chiralities investigated exhibited similar trends for both phospholipid-based agents (Supplementary Figures S17C, S18C, S19C). Time-course measurements up to 180 minutes demonstrated relative stability during this period (Supplementary Figures S20, S21, S22).

It is likely that proteins, such as BSA, NFDM, and casein, may cause specific changes by forming a homogenous protein corona. These proteins may interact with the nanotube surface through hydrophobic adsorption and/or charge-charge interactions with the ssDNA. Similarly, the interactions between cationic polymers such as PEI and PLK with SWCNT surfaces may be aided by strong columbic interactions, allowing a robust and stable corona to form. Further, we propose that the cationic charge and linear structure of PLK may aid in strong negative columbic interactions with SWCNT-(TAT)_6_ without flocculation as observed for cationic but branched PEI. The PEG construct used is anionic and therefore expected to show little interaction due to charge repulsion with the ssDNA. Further, changes in absorbance due to interaction with 16:0 PE2000PEG are likely due to the hydrophobic interaction of the lipid tails with the SWCNT surface^78^. DSPE-PEG (NH2) is a zwitterionic PEG modified lipid with a long hydrophobic tail^78^, which may facilitate hydrophobic and/or charge-charge interactions. However, for screening of serum interference, this phospholipid did not exhibit sufficient passivation.

### Construction and validation of an IL-6 nanosensor

We then sought to apply the passivation tools developed here for use with a clinically-relevant nanosensor device. We prepared and characterized a nanosensor comprised of a monoclonal IL-6 antibody conjugated to SWCNT-(TAT)_6_-NH2 (Supplementary Figure S23). Dynamic light scattering (DLS) revealed successful conjugation as the correlation coefficient demonstrated a larger relative particle size for the conjugate nanosensor compared to unconjugated SWCNT-(TAT)_6_-NH2 complexes as previously described (Supplementary Figure S23A)^4, 5^. Further, electrophoretic light scattering demonstrated a more negative ζ-potential for SWCNT-(TAT)_6_-NH2 than that for the IL-6 nanosensor conjugate, confirming successful antibody conjugation as in prior works (Supplementary Figure S23B)^4, 5^.

We first tested the sensor fluorescence response to 5,250 ng/mL IL-6 in simple buffer conditions of 1x PBS. The center wavelength of the (7,5) chirality of the nanosensor blue shifted 3.6 ± 0.63 nm (p = 0.13), though this response was not significant (Supplementary Figure S23C). Next, we tested the response of the 50x BSA passivated IL-6 nanosensor to the same concentration of IL-6 in 10% FBS. We saw minimal shifting of the (7,5) chirality of 0.43 ± 0.29 nm (p = 0.09) (Supplementary Figure S23D).

### Detection of IL-6 in human serum

Having validated basic IL-6 nanosensor function, we next sought to evaluate its function in a simulated clinical environment using human serum. To do so, we passivated separate batches of IL-6 nanosensor with BSA, PLK, and NFDM at a ratio of 50x. These were chosen as they exhibited substantial screening responses above, as well as the formation of a stable homogenous corona as revealed by absorbance spectroscopy. Serum IL-6 levels in healthy patients typically range from 1-10 pg/mL^34, 38^. For patients with health conditions such COVID-19 or sepsis, serum concentrations are elevated, reaching 10,000 pg/mL or greater^38, 79, 80^. Considering this, we simulated disease conditions by spiking known quantities of IL-6 into human serum at concentrations of 25; 250; 2,500; and 25,000 pg/mL. We then compared shifts in fluorescence center wavelength for each chirality and determined whether the shifts were statistically significant and consistent compared to controls.

We found that, by evaluating four chiralities at once, we were able to positively identify each concentration investigated (**Figure 6**), including the extremely low, and clinically relevant, 25 pg/mL. Following PLK passivation, we observed that the (7,5) and (8,7) chiralities demonstrated significant shifts in the presence of 25 pg/mL IL-6 (Supplementary Figures S24, S25) and the (7,6) demonstrated a significant shift in the presence of 250 pg/mL (Supplementary Figure S26). All four chiralities that we observed exhibited significant shifts in the presence of 2,500 pg/mL (Supplementary Figures S24 – S27) while the (8,7) and (9,4) demonstrated significant shifts in the presence of 25,000 pg/mL (Supplementary Figures S25, S27). Of the other passivation agents investigated (BSA and NFDM), only the BSA-passivated nanosensor demonstrated any statistically significant shifts, specifically the (9,4) chirality with 250 pg/mL IL-6 (Supplementary Figure 27). This demonstrated multichiral nanosensor with highly sensitive and quantitative IL-6 detection in human serum gives rise to the possibility of rapid detection at home or the patient bedside.

**Figure 6.**
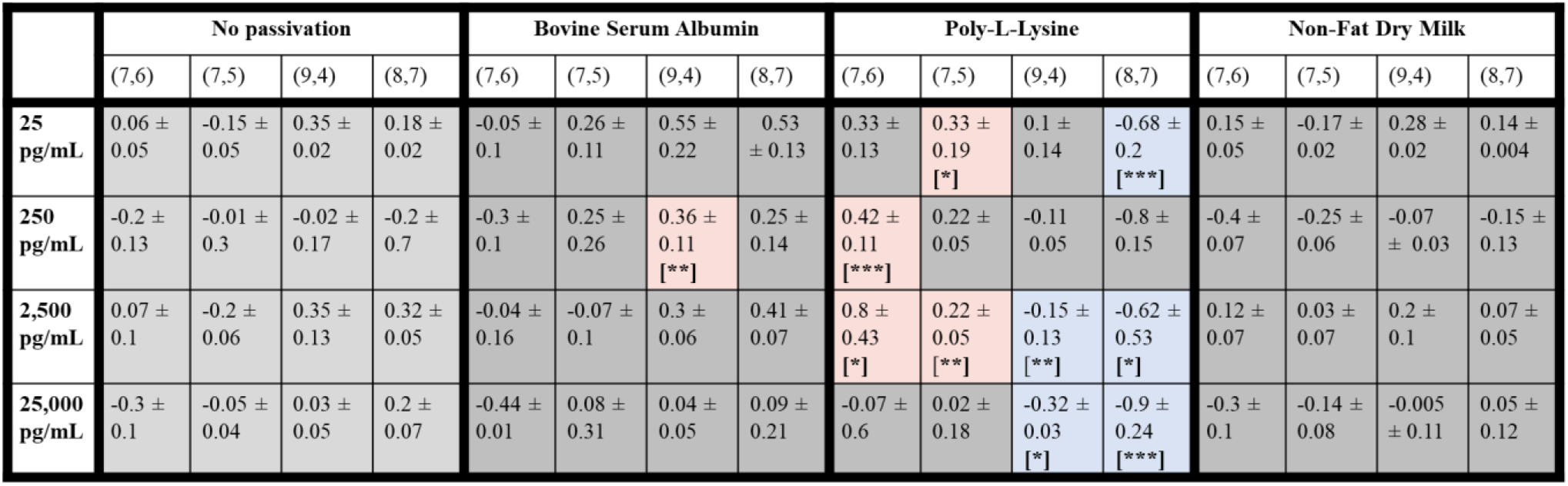
Detection of IL-6 in human serum. Shifts in center wavelength (mean ± standard deviation) are indicated for each of the (7,6), (7,5), (9,4), and (8,7) SWCNT chiralities after addition of the indicated concentration of IL-6, categorized by passivation agent (note: a “-” indicates blue shift). Light grey indicates non-passivated controls and dark grey indicates changes which are not statistically significant. Blue indicates statistically significant blue shifts, while red indicates statistically significant red shifts. * = p < 0.05; ** = p < 0.01; *** = p < 0.001).

Based on a substantial literature search, this work represents a dramatic increase in the sensitivity of SWCNT-based optical sensors in human serum, and the first such for IL-6 inflammatory cytokines. In prior studies, antibody-conjugated SWCNT-based optical nanosensors for cancer biomarkers have shown a limit of detection (LOD) of 33,800 pg/mL (2.6 nM) for the ovarian cancer biomarker HE-4 in human serum^4^. Here, we demonstrated that the PLK passivated nanosensor detects IL-6 concentrations as low as 25 pg/mL in human serum, 1,352 times lower than that which was previously reported. It also provides detection across four orders of magnitude, from 25 pg/mL to 25,000 pg/mL^4^. IL-6 levels in serum are reported to be above 20 pg/mL in patients diagnosed with cancer, neurological diseases, sepsis, and COVID-19^38^.

Concentration levels above 500 pg/mL and 7,500 pg/mL correlate with patient mortality in 11% and 37% of sepsis cases, respectively^81^. The IL-6 nanosensor presented here demonstrates detection, in clinically-relevant serum conditions, well within these diagnostic concentration ranges. In addition to the clearly-established bedside diagnostic potential of this nanosensor construct, combining this with the established in vivo detection applicability of SWCNTs^35, 41^ portends a strong potential for implantable diagnostic device design.

We also found that the ability to detect different IL-6 concentrations differs across nanotube chiralities for a given passivation agent. Many factors influence the success of passivation—size, conformation, hydrophobicity, and ionic charge of a given passivation agent, as well as the atomic composition, surface roughness, and curvature of the nanomaterial being passivated, plus the pH, temperature, and ionic strength of the local solute environment^82^. It is known that SWCNT chiralities differ in surface composition and size due to varying ssDNA wrapping abilities^83-86^. In this study, the charge-charge interactions of PLK with ssDNA are expected to dominate the passivation process (**Figure 7**)^71^. Different ssDNA sequences are known to provide varying degrees of surface coverage for specific SWCNT chiralities, which could potentially affect the extent of PLK passivation success across different chiralities^10, 84-86^. We propose these variations contribute to variations in functionality for a given SWCNT chirality.

**Figure 7.**
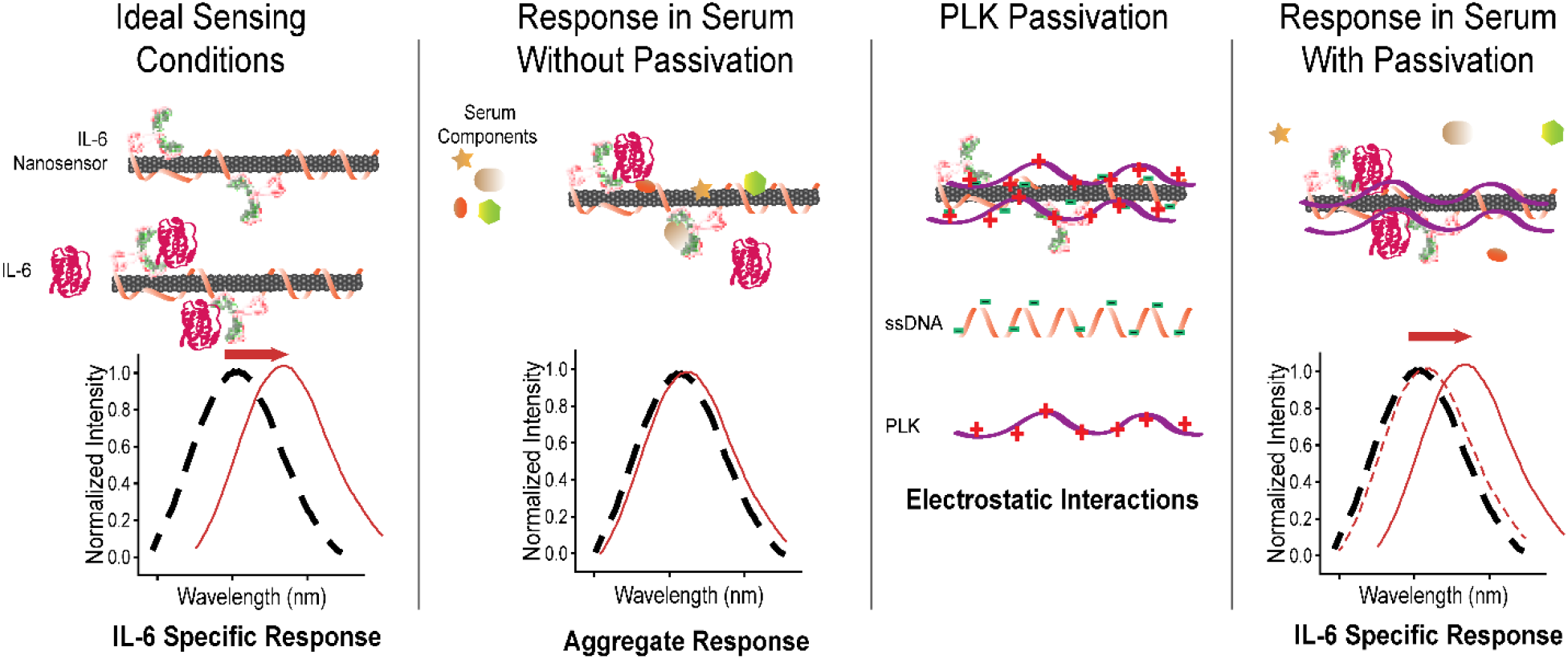
Diagram of proposed mechanism of action for PLK-induced sensing of IL-6 in human serum.

## Conclusions

In this work, we observed highly sensitive detection of the clinically-relevant cytokine IL-6 in human serum across a broad functional range of concentrations. To achieve this, we first screened three classes of biomolecules—proteins, polymers, and amphiphiles—as potential passivating agents. We found that bovine serum albumin (BSA), non-fat dry milk (NFDM), and casein – all proteins, and poly-L-lysine (PLK), a polymer, were the most successful passivation agents in FBS screening. We found that 50x and 100x mass ratios of passivating agents to SWCNTs were more successful than 5x and 25x mass ratios. These results were corroborated by absorption spectroscopy, demonstrating a stable surface coverage for each. With PLK and/or BSA passivation, an engineered IL-6 nanosensor demonstrated clinically-relevant detection as low as 25 pg/mL, ranging up to 25,000 pg/mL upon observation of multiple nanotube chiralities. We expect this study to provide rational strategies to screen interference from heterogeneously-formed coronas upon introduction to complex sensing conditions, improving the selectivity and sensitivity of, specifically, SWCNT-based optical nanosensors, and more broadly, other nanosensors, for clinical applications.

## Acknowledgements

The authors wish to acknowledge all members of the Williams Lab for discussion and feedback. This work was supported by NIH R35GM142833 (RMW) and the support of The City College of New York Grove School of Engineering. It was also supported by a Dissertation Year Fellowship to PG from the CUNY Graduate Center.

## Notes

### Competing Interest Statement

The authors have declared no competing interest.

## References

1. Aravind Kumar, J.; Krithiga, T.; Venkatesan, D., Carbon Nanotubes: Synthesis, Properties and Applications. In 21st Century Surface Science, Phuong, P.; Pratibha, G.; Samir, K.; Kavita, Y., Eds. IntechOpen: Rijeka, 2020; p Ch. 2.

2. Hendler-Neumark, A.; Bisker, G., Fluorescent Single-Walled Carbon Nanotubes for Protein Detection. Sensors (Basel) 2019, 19 (24).

3. Boghossian, A. A.; Zhang, J.; Barone, P. W.; Reuel, N. F.; Kim, J.-H.; Heller, D. A.; Ahn, J.-H.; Hilmer, A. J.; Rwei, A.; Arkalgud, J. R.; Zhang, C. T.; Strano, M. S., Near-Infrared Fluorescent Sensors based on Single-Walled Carbon Nanotubes for Life Sciences Applications. ChemSusChem 2011, 4 (7), 848–863.

4. Williams, R. M.; Lee, C.; Galassi, T. V.; Harvey, J. D.; Leicher, R.; Sirenko, M.; Dorso, M. A.; Shah, J.; Olvera, N.; Dao, F.; Levine, D. A.; Heller, D. A., Noninvasive ovarian cancer biomarker detection via an optical nanosensor implant. Sci Adv 2018, 4 (4), eaaq1090.

5. Williams, R. M.; Lee, C.; Heller, D. A., A Fluorescent Carbon Nanotube Sensor Detects the Metastatic Prostate Cancer Biomarker uPA. ACS Sens 2018, 3 (9), 1838–1845.

6. Harvey, J. D.; Williams, R. M.; Tully, K. M.; Baker, H. A.; Shamay, Y.; Heller, D. A., An in Vivo Nanosensor Measures Compartmental Doxorubicin Exposure. Nano Letters 2019, 19 (7), 4343–4354.

7. Kosuge, H.; Sherlock, S. P.; Kitagawa, T.; Dash, R.; Robinson, J. T.; Dai, H.; McConnell, M. V., Near Infrared Imaging and Photothermal Ablation of Vascular Inflammation Using Single-Walled Carbon Nanotubes. Journal of the American Heart Association 2012, 1 (6), e002568.

8. Elizarova, S.; Chouaib, A. A.; Shaib, A.; Hill, B.; Mann, F.; Brose, N.; Kruss, S.; Daniel, J. A., A fluorescent nanosensor paint detects dopamine release at axonal varicosities with high spatiotemporal resolution. Proceedings of the National Academy of Sciences 2022, 119 (22), e2202842119.

9. Bachilo, S. M.; Strano, M. S.; Kittrell, C.; Hauge, R. H.; Smalley, R. E.; Weisman, R. B., Structure-Assigned Optical Spectra of Single-Walled Carbon Nanotubes. Science 2002, 298 (5602), 2361–2366.

10. Tu, X.; Zheng, M., A DNA-based approach to the carbon nanotube sorting problem. Nano Research 2008, 1 (3), 185–194.

11. Streit, J. K.; Bachilo, S. M.; Ghosh, S.; Lin, C.-W.; Weisman, R. B., Directly Measured Optical Absorption Cross Sections for Structure-Selected Single-Walled Carbon Nanotubes. Nano Letters 2014, 14 (3), 1530–1536.

12. Ackermann, J.; Metternich, J. T.; Herbertz, S.; Kruss, S., Biosensing with Fluorescent Carbon Nanotubes. Angewandte Chemie International Edition 2022, 61 (18), e202112372.

13. Jeong, S.; Yang, D.; Beyene, A. G.; Del Bonis-O’Donnell, J. T.; Gest, A. M. M.; Navarro, N.; Sun, X.; Landry, M. P., High-throughput evolution of near-infrared serotonin nanosensors. Science Advances 2019, 5 (12), eaay3771.

14. Pinals, R. L.; Ledesma, F.; Yang, D.; Navarro, N.; Jeong, S.; Pak, J. E.; Kuo, L.; Chuang, Y.-C.; Cheng, Y.-W.; Sun, H.-Y.; Landry, M. P., Rapid SARS-CoV-2 Spike Protein Detection by Carbon Nanotube-Based Near-Infrared Nanosensors. Nano Letters 2021, 21 (5), 2272–2280.

15. Kim, M.; Chen, C.; Wang, P.; Mulvey, J. J.; Yang, Y.; Wun, C.; Antman-Passig, M.; Luo, H.-B.; Cho, S.; Long-Roche, K.; Ramanathan, L. V.; Jagota, A.; Zheng, M.; Wang, Y.; Heller, D. A., Detection of ovarian cancer via the spectral fingerprinting of quantum-defect-modified carbon nanotubes in serum by machine learning. Nature Biomedical Engineering 2022, 6 (3), 267–275.

16. Budhathoki-Uprety, J.; Shah, J.; Korsen, J. A.; Wayne, A. E.; Galassi, T. V.; Cohen, J. R.; Harvey, J. D.; Jena, P. V.; Ramanathan, L. V.; Jaimes, E. A.; Heller, D. A., Synthetic molecular recognition nanosensor paint for microalbuminuria. Nature Communications 2019, 10 (1), 3605.

17. Loewenthal, D.; Kamber, D.; Bisker, G., Monitoring the Activity and Inhibition of Cholinesterase Enzymes using Single-Walled Carbon Nanotube Fluorescent Sensors. Analytical Chemistry 2022, 94 (41), 14223–14231.

18. Albarghouthi, F. M.; Williams, N. X.; Doherty, J. L.; Lu, S.; Franklin, A. D., Passivation Strategies for Enhancing Solution-Gated Carbon Nanotube Field-Effect Transistor Biosensing Performance and Stability in Ionic Solutions. ACS Applied Nano Materials 2022, 5 (10), 15865–15874.

19. Zhang, J.; Kruss, S.; Hilmer, A. J.; Shimizu, S.; Schmois, Z.; De La Cruz, F.; Barone, P. W.; Reuel, N. F.; Heller, D. A.; Strano, M. S., A Rapid, Direct, Quantitative, and Label-Free Detector of Cardiac Biomarker Troponin T Using Near-Infrared Fluorescent Single-Walled Carbon Nanotube Sensors. Advanced Healthcare Materials 2014, 3 (3), 412–423.

20. Hu, D.; Yang, L.; Deng, S.; Hao, Y.; Zhang, K.; Wang, X.; Liu, Y.; Liu, H.; Chen, Y.; Xie, M., Development of nanosensor by bioorthogonal reaction for multi-detection of the biomarkers of hepatocellular carcinoma. Sensors and Actuators B: Chemical 2021, 334, 129653.

21. Rezaee, M.; Behnam, B.; Banach, M.; Sahebkar, A., The Yin and Yang of carbon nanomaterials in atherosclerosis. Biotechnology Advances 2018, 36 (8), 2232–2247.

22. Sardesai, N. P.; Barron, J. C.; Rusling, J. F., Carbon Nanotube Microwell Array for Sensitive Electrochemiluminescent Detection of Cancer Biomarker Proteins. Analytical Chemistry 2011, 83 (17), 6698–6703.

23. Shao, N.; Wickstrom, E.; Panchapakesan, B., Nanotube-antibody biosensor arrays for the detection of circulating breast cancer cells. Nanotechnology 2008, 19 (46), 465101.

24. Khosravi Ardakani, H.; Gerami, M.; Chashmpoosh, M.; Omidifar, N.; Gholami, A., Recent Progress in Nanobiosensors for Precise Detection of Blood Glucose Level. Biochemistry Research International 2022, 2022, 2964705.

25. Taguchi, M.; Ptitsyn, A.; McLamore, E. S.; Claussen, J. C., Nanomaterial-mediated biosensors for monitoring glucose. Journal of Diabetes Science and Technology 2014, 8 (2), 403–411.

26. Li, T.; Soelberg, S. D.; Taylor, Z.; Sakthivelpathi, V.; Furlong, C. E.; Kim, J.-H.; Ahn, S.-g.; Han, P. D.; Starita, L. M.; Zhu, J.; Chung, J.-H., Highly Sensitive Immunoresistive Sensor for Point-Of-Care Screening for COVID-19. Biosensors 2022, 12 (3), 149.

27. Cash, K. J.; Clark, H. A., In Vivo Histamine Optical Nanosensors. Sensors 2012, 12 (9), 11922–11932.

28. Iverson, N. M.; Barone, P. W.; Shandell, M.; Trudel, L. J.; Sen, S.; Sen, F.; Ivanov, V.; Atolia, E.; Farias, E.; McNicholas, T. P.; Reuel, N.; Parry, N. M. A.; Wogan, G. N.; Strano, M. S., In vivo biosensing via tissue-localizable near-infrared-fluorescent single-walled carbon nanotubes. Nature Nanotechnology 2013, 8 (11), 873–880.

29. Hofferber, E.; Meier, J.; Herrera, N.; Stapleton, J.; Calkins, C.; Iverson, N., Detection of single walled carbon nanotube based sensors in a large mammal. Nanomedicine: Nanotechnology, Biology and Medicine 2022, 40, 102489.

30. Galassi, T. V.; Jena, P. V.; Shah, J.; Ao, G.; Molitor, E.; Bram, Y.; Frankel, A.; Park, J.; Jessurun, J.; Ory, D. S., An optical nanoreporter of endolysosomal lipid accumulation reveals enduring effects of diet on hepatic macrophages in vivo. Science translational medicine 2018, 10 (461), eaar2680.

31. Harvey, J. D.; Jena, P. V.; Baker, H. A.; Zerze, G. H.; Williams, R. M.; Galassi, T. V.; Roxbury, D.; Mittal, J.; Heller, D. A., A carbon nanotube reporter of microRNA hybridization events in vivo. Nature Biomedical Engineering 2017, 1, 0041.

32. Miri, Y.; Leander, K.; Eriksson, P.; Gigante, B.; Ziegler, L., Interleukin 6 trans-signalling and the risk of future cardiovascular events in men and women. Open Heart 2021, 8 (2), e001694.

33. Wainstein, M. V.; Mossmann, M.; Araujo, G. N.; Gonçalves, S. C.; Gravina, G. L.; Sangalli, M.; Veadrigo, F.; Matte, R.; Reich, R.; Costa, F. G.; Andrades, M.; da Silva, A. M. V.; Bertoluci, M. C., Elevated serum interleukin-6 is predictive of coronary artery disease in intermediate risk overweight patients referred for coronary angiography. Diabetology & Metabolic Syndrome 2017, 9 (1), 67.

34. Villar-Fincheira, P.; Sanhueza-Olivares, F.; Norambuena-Soto, I.; Cancino-Arenas, N.; Hernandez-Vargas, F.; Troncoso, R.; Gabrielli, L.; Chiong, M., Role of Interleukin-6 in Vascular Health and Disease. Frontiers in Molecular Biosciences 2021, 8.

35. Potere, N.; Batticciotto, A.; Vecchié, A.; Porreca, E.; Cappelli, A.; Abbate, A.; Dentali, F.; Bonaventura, A., The role of IL-6 and IL-6 blockade in COVID-19. Expert review of clinical immunology 2021, 17 (6), 601–618.

36. Chonov, D. C.; Ignatova, M. M. K.; Ananiev, J. R.; Gulubova, M. V., IL-6 Activities in the Tumour Microenvironment. Part 1. Open Access Maced J Med Sci 2019, 7 (14), 2391–2398.

37. Kumari, N.; Dwarakanath, B. S.; Das, A.; Bhatt, A. N., Role of interleukin-6 in cancer progression and therapeutic resistance. Tumour Biol 2016, 37 (9), 11553–11572.

38. McCrae, L. E.; Ting, W.-T.; Howlader, M. M. R., Advancing electrochemical biosensors for interleukin-6 detection. Biosensors and Bioelectronics: X 2023, 13, 100288.

39. Tanaka, T.; Narazaki, M.; Kishimoto, T., IL-6 in inflammation, immunity, and disease. Cold Spring Harb Perspect Biol 2014, 6 (10), a016295.

40. Hirano, T., IL-6 in inflammation, autoimmunity and cancer. International Immunology 2020, 3 (3), 127–148.

41. Yoshida, Y.; Tanaka, T., Interleukin 6 and rheumatoid arthritis. BioMed research international 2014, 2014.

42. Wognum, A. W.; van Gils, F. C. J. M.; Wagemaker, G., Flow Cytometric Detection of Receptors for Interleukin-6 on Bone Marrow and Peripheral Blood Cells of Humans and Rhesus Monkeys. Blood 1993, 81 (8), 2036–2043.

43. Grellner, W., Time-dependent immunohistochemical detection of proinflammatory cytokines (IL-1β, IL-6, TNF-α) in human skin wounds. Forensic Science International 2002, 130 (2), 90–96.

44. Luo, J.; Gopinath, S. C. B.; Subramaniam, S.; Wu, Z., Arthritis biosensing: Aptamer-antibody-mediated identification of biomarkers by ELISA. Process Biochemistry 2022, 121, 396–402.

45. Martin, K.; Viera, K.; Petr, C.; Marie, N.; Eva, T., Simultaneous analysis of cytokines and costimulatory molecules concentrations by ELISA technique andof probabilities of measurable concentrations of interleukins IL-2, IL-4, IL-5, IL-6, CXCL8 (IL-8), IL-10, IL-13 occurring in plasma of healthy blood donors. Mediators of Inflammation 2006, 2006, 065237.

46. Kaur, J.; Preethi, M.; Srivastava, R.; Borse, V., Role of IL-6 and IL-8 biomarkers for optical and electrochemical based point-of-care detection of oral cancer. Biosensors and Bioelectronics: X 2022, 11, 100212.

47. Rahbar, M.; Wu, Y.; Subramony, J. A.; Liu, G., Sensitive Colorimetric Detection of Interleukin-6 via Lateral Flow Assay Incorporated Silver Amplification Method. Frontiers in Bioengineering and Biotechnology 2021, 9.

48. Hao, Z.; Pan, Y.; Shao, W.; Lin, Q.; Zhao, X., Graphene-based fully integrated portable nanosensing system for on-line detection of cytokine biomarkers in saliva. Biosensors and Bioelectronics 2019, 134, 16–23.

49. Adrover-Jaume, C.; Alba-Patiño, A.; Clemente, A.; Santopolo, G.; Vaquer, A.; Russell, S. M.; Barón, E.; González del Campo, M. d. M.; Ferrer, J. M.; Berman-Riu, M.; García-Gasalla, M.; Aranda, M.; Borges, M.; de la Rica, R., Paper biosensors for detecting elevated IL-6 levels in blood and respiratory samples from COVID-19 patients. Sensors and Actuators B: Chemical 2021, 330, 129333.

50. Saatçi, E.; Natarajan, S., State-of-the-art colloidal particles and unique interfaces-based SARS-CoV-2 detection methods and COVID-19 diagnosis. Current Opinion in Colloid & Interface Science 2021, 55, 101469.

51. Hu, W.-P.; Wu, Y.-M.; Vu, C.-A.; Chen, W.-Y., Ultrasensitive Detection of Interleukin 6 by Using Silicon Nanowire Field-Effect Transistors. Sensors 2023, 23 (2), 625.

52. Nguyen, V. H.; Lee, B. J., Protein corona: a new approach for nanomedicine design. Int J Nanomedicine 2017, 12, 3137–3151.

53. Pinals, R. L.; Yang, D.; Rosenberg, D. J.; Chaudhary, T.; Crothers, A. R.; Iavarone, A. T.; Hammel, M.; Landry, M. P., Quantitative Protein Corona Composition and Dynamics on Carbon Nanotubes in Biological Environments. Angew Chem Int Ed Engl 2020, 59 (52), 23668–23677.

54. Iverson, N. M.; Bisker, G.; Farias, E.; Ivanov, V.; Ahn, J.; Wogan, G. N.; Strano, M. S., Quantitative tissue spectroscopy of near infrared fluorescent nanosensor implants. Journal of biomedical nanotechnology 2016, 12 (5), 1035–1047.

55. Monopoli, M. P.; Aberg, C.; Salvati, A.; Dawson, K. A., Biomolecular coronas provide the biological identity of nanosized materials. Nat Nanotechnol 2012, 7 (12), 779–86.

56. Nel, A. E.; Mädler, L.; Velegol, D.; Xia, T.; Hoek, E. M. V.; Somasundaran, P.; Klaessig, F.; Castranova, V.; Thompson, M., Understanding biophysicochemical interactions at the nano–bio interface. Nature Materials 2009, 8 (7), 543–557.

57. Boghossian, A. A.; Zhang, J.; Barone, P. W.; Reuel, N. F.; Kim, J. H.; Heller, D. A.; Ahn, J. H.; Hilmer, A. J.; Rwei, A.; Arkalgud, J. R.; Zhang, C. T.; Strano, M. S., Near-infrared fluorescent sensors based on single-walled carbon nanotubes for life sciences applications. ChemSusChem 2011, 4 (7), 848–63.

58. Mahmood, T.; Yang, P. C., Western blot: technique, theory, and trouble shooting. N Am J Med Sci 2012, 4 (9), 429–34.

59. Jiang, X.; Wu, M.; Albo, J.; Rao, Q., Non-Specific Binding and Cross-Reaction of ELISA: A Case Study of Porcine Hemoglobin Detection. Foods 2021, 10 (8), 1708.

60. Mehdi, F.; Chattopadhyay, S.; Thiruvengadam, R.; Yadav, S.; Kumar, M.; Sinha, S. K.; Goswami, S.; Kshetrapal, P.; Wadhwa, N.; Chandramouli Natchu, U.; Sopory, S.; Koundinya Desiraju, B.; Pandey, A. K.; Das, A.; Verma, N.; Sharma, N.; Sharma, P.; Bhartia, V.; Gosain, M.; Lodha, R.; Lamminmäki, U.; Shrivastava, T.; Bhatnagar, S.; Batra, G., Development of a Fast SARS-CoV-2 IgG ELISA, Based on Receptor-Binding Domain, and Its Comparative Evaluation Using Temporally Segregated Samples From RT-PCR Positive Individuals. Front Microbiol 2020, 11, 618097.

61. Ma, G. J.; Ferhan, A. R.; Jackman, J. A.; Cho, N.-J., Conformational flexibility of fatty acid-free bovine serum albumin proteins enables superior antifouling coatings. Communications Materials 2020, 1 (1), 45.

62. Lim, C. S.; Krishnan, G.; Sam, C. K.; Ng, C. C., On optimizing the blocking step of indirect enzyme-linked immunosorbent assay for Epstein-Barr virus serology. Clinica Chimica Acta 2013, 415, 158–161.

63. Yang, D.; Yang, S. J.; Del Bonis-O’Donnell, J. T.; Pinals, R. L.; Landry, M. P., Mitigation of Carbon Nanotube Neurosensor Induced Transcriptomic and Morphological Changes in Mouse Microglia with Surface Passivation. ACS Nano 2020, 14 (10), 13794–13805.

64. Jeyachandran, Y. L.; Mielczarski, J. A.; Mielczarski, E.; Rai, B., Efficiency of blocking of nonspecific interaction of different proteins by BSA adsorbed on hydrophobic and hydrophilic surfaces. J Colloid Interface Sci 2010, 341 (1), 136–42.

65. Reimhult, K.; Petersson, K.; Krozer, A., QCM-D Analysis of the Performance of Blocking Agents on Gold and Polystyrene Surfaces. Langmuir 2008, 24 (16), 8695–8700.

66. Hadidi, N.; Shahbahrami Moghadam, N.; Pazuki, G.; Parvin, P.; Shahi, F., In Vitro Evaluation of DSPE-PEG (5000) Amine SWCNT Toxicity and Efficacy as a Novel Nanovector Candidate in Photothermal Therapy by Response Surface Methodology (RSM). Cells 2021, 10 (11), 2874.

67. Sacchetti, C.; Rapini, N.; Magrini, A.; Cirelli, E.; Bellucci, S.; Mattei, M.; Rosato, N.; Bottini, N.; Bottini, M., In Vivo Targeting of Intratumor Regulatory T Cells Using PEG-Modified Single-Walled Carbon Nanotubes. Bioconjugate Chemistry 2013, 24 (6), 852–858.

68. Campagnolo, L.; Massimiani, M.; Palmieri, G.; Bernardini, R.; Sacchetti, C.; Bergamaschi, A.; Vecchione, L.; Magrini, A.; Bottini, M.; Pietroiusti, A., Biodistribution and toxicity of pegylated single wall carbon nanotubes in pregnant mice. Particle and Fibre Toxicology 2013, 10 (1), 21.

69. Viswanathan, S.; Rani, C.; Vijay Anand, A.; Ho, J.-a. A., Disposable electrochemical immunosensor for carcinoembryonic antigen using ferrocene liposomes and MWCNT screen-printed electrode. Biosensors and Bioelectronics 2009, 24 (7), 1984–1989.

70. Bilalis, P.; Katsigiannopoulos, D.; Avgeropoulos, A.; Sakellariou, G., Non-covalent functionalization of carbon nanotubes with polymers. RSC Advances 2014, 4 (6), 2911–2934.

71. Zheng, M.; Pan, M.; Zhang, W.; Lin, H.; Wu, S.; Lu, C.; Tang, S.; Liu, D.; Cai, J., Poly(α-l-lysine)-based nanomaterials for versatile biomedical applications: Current advances and perspectives. Bioactive Materials 2021, 6 (7), 1878–1909.

72. Harvey, J. D.; Jena, P. V.; Baker, H. A.; Zerze, G. H.; Williams, R. M.; Galassi, T. V.; Roxbury, D.; Mittal, J.; Heller, D. A., A Carbon Nanotube Reporter of miRNA Hybridization Events In Vivo. Nat Biomed Eng 2017, 1.

73. Hong, X.; Meng, Y.; Kalkanis, S. N., Serum proteins are extracted along with monolayer cells in plasticware and interfere with protein analysis. J Biol Methods 2016, 3 (4).

74. Wilbanks, D. J.; Lee, M. R.; Rahimi, Y. S.; Lucey, J. A., Comparison of micellar casein isolate and nonfat dry milk for use in the production of high-protein cultured milk products. Journal of Dairy Science 2023, 106 (1), 61–74.

75. Streit, J. K.; Bachilo, S. M.; Ghosh, S.; Lin, C. W.; Weisman, R. B., Directly measured optical absorption cross sections for structure-selected single-walled carbon nanotubes. Nano Lett 2014, 14 (3), 1530–6.

76. Weisman, R. B.; Bachilo, S. M., Dependence of Optical Transition Energies on Structure for Single-Walled Carbon Nanotubes in Aqueous Suspension: An Empirical Kataura Plot. Nano Letters 2003, 3 (9), 1235–1238.

77. Galassi, T.; Jena, P.; Roxbury, D.; Heller, D., Single Nanotube Spectral Imaging To Determine Molar Concentrations of Isolated Carbon Nanotube Species. Analytical Chemistry 2017, 89.

78. Webb, M. S.; Saxon, D.; Wong, F. M. P.; Lim, H. J.; Wang, Z.; Bally, M. B.; Choi, L. S. L.; Cullis, P. R.; Mayer, L. D., Comparison of different hydrophobic anchors conjugated to poly(ethylene glycol): effects on the pharmacokinetics of liposomal vincristine. Biochimica et Biophysica Acta (BBA) - Biomembranes 1998, 1372 (2), 272–282.

79. Strand, V.; Boklage, S. H.; Kimura, T.; Joly, F.; Boyapati, A.; Msihid, J., High levels of interleukin-6 in patients with rheumatoid arthritis are associated with greater improvements in healthrelated quality of life for sarilumab compared with adalimumab. Arthritis Research & Therapy 2020, 22 (1), 250.

80. Sun, H.; Guo, P.; Zhang, L.; Wang, F., Serum Interleukin-6 Concentrations and the Severity of COVID-19 Pneumonia: A Retrospective Study at a Single Center in Bengbu City, Anhui Province, China, in January and February 2020. Med Sci Monit 2020, 26, e926941.

81. Russell, C.; Ward, A. C.; Vezza, V.; Hoskisson, P.; Alcorn, D.; Steenson, D. P.; Corrigan, D. K., Development of a needle shaped microelectrode for electrochemical detection of the sepsis biomarker interleukin-6 (IL-6) in real time. Biosensors and Bioelectronics 2019, 126, 806–814.

82. Park, J. H.; Sut, T. N.; Jackman, J. A.; Ferhan, A. R.; Yoon, B. K.; Cho, N.-J., Controlling adsorption and passivation properties of bovine serum albumin on silica surfaces by ionic strength modulation and cross-linking. Physical Chemistry Chemical Physics 2017, 19 (13), 8854–8865.

83. Gu, Z.; Yang, Z.; Chong, Y.; Ge, C.; Weber, J. K.; Bell, D. R.; Zhou, R., Surface Curvature Relation to Protein Adsorption for Carbon-based Nanomaterials. Scientific Reports 2015, 5 (1), 10886.

84. Hinkle, K. R., Molecular dynamics simulations reveal single-stranded DNA (ssDNA) forms ordered structures upon adsorbing onto single-walled carbon nanotubes (SWCNT). Colloids Surf B Biointerfaces 2022, 212, 112343.

85. Neihsial, S.; Periyasamy, G.; Samanta, P. K.; Pati, S. K., Understanding the Binding Mechanism of Various Chiral SWCNTs and ssDNA: A Computational Study. The Journal of Physical Chemistry B 2012, 116 (51), 14754–14759.

86. Zheng, Y.; Alizadehmojarad, A. A.; Bachilo, S. M.; Kolomeisky, A. B.; Weisman, R. B., Dye Quenching of Carbon Nanotube Fluorescence Reveals Structure-Selective Coating Coverage. ACS Nano 2020, 14 (9), 12148–12158.

