## Supplementary Information for "Optical nanosensor passivation enables highly sensitive detection of the inflammatory cytokine IL-6"

**Supplementary Calculations**

Per manufacturer specifications (NanoIntegris Technologies, Inc), the average diameter for SWCNT ranges between 0.8 nm-1.2 nm. Assuming an average diameter of 1.0 nm, the circumference of an individual SWCNT is 3.14 nm using the equation $C = 2 \pi r$. Each C-C bond is 1.54 angstroms long. Hence, we estimate that, along the circumference, there are (31.4/1.54) angstroms = about 20 carbon atoms. Per the manufacturer, for every 0.283 nm length, there are 4 x 20 = 80 carbon atoms. The atomic mass of single carbon atom is 12.01 amu.

The previously reported average length of similarly-prepared SWCNT is 166 nm[1], therefore the average mass of an individual SWCNT is (166/0.283) x 80 x 12.01 = 563,578.8 amu = 563.5 kDa. We used 0.5 mg/L SWCNT-$(TAT)_6$ for all our experiments, and using the calculated mass per SWCNT derives a concentration of 0.887 nM.

For 50x mass ratio passivation, 0.5 mg/L x 50 = 25 mg/L concentration is used for each passivation agent. We calculated molar concentration used for each passivation agent as (25 mg/L)/(molecular weight) = molarity.

The molecular weights reported in the literature for BSA[2] and casein[3] are 66.4 kDa and 20-25 kDa, respectively. NFDM is a heterogenous mixture and therefore no molecular weight is available. We calculated that a 50x mass ratio is 376.5 nM and 1.25 uM-1uM for BSA and casein, respectively. Similarly, for PLK (molecular weight 70,000-150,000 Da, per the manufacturer; Advanced Biomatrix), PEI (molecular weight 10,000 Da, per the manufacturer; Alfa-Aesar), and PEG (M.W. 1500 Da, per the manufacturer; Sigma Aldrich), 25 mg/L is 166.7 nM-357.1 nM, 2500 nM, and 16,670 nM respectively. For DSPE PEG (NH2) (molecular weight 2000 g, per the manufacturer; BroadPharm) and 16:0 PE PEG (formula weight 2749.42 g, per the manufacturer; Avanti Polar Lipids Inc), it is 12,500 nM and 9093 nM, respectively.

Therefore, the estimated molar ratio (as opposed to an exact 50x mass ratio reported in the main text), for each passivation agent compared to SWCNT-$(TAT)_6$ was: 425:1 for BSA, 1409-1127:1 for casein, 188-403:1 for PLK, 2819:1 for PEI, 1.2E4:1 for PEG, 1.409E4:1 for DSPE PEG (NH2), and 1.025E4:1 for 16:0 PE PEG.

### Supplementary Figures

| Class | Structure |
|---|---|
| Polymers | • Polyethylene Glycol 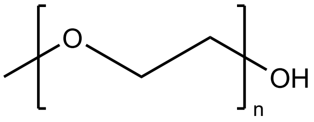 |
| | • Polyethylene Imine 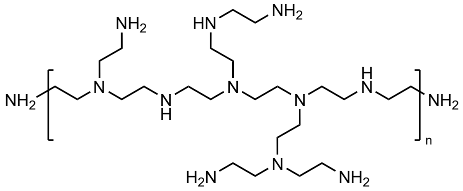 |
| | • Poly-L-Lysine 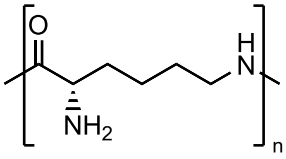 |
| Surfactants | • 16:0 PE2000PEG 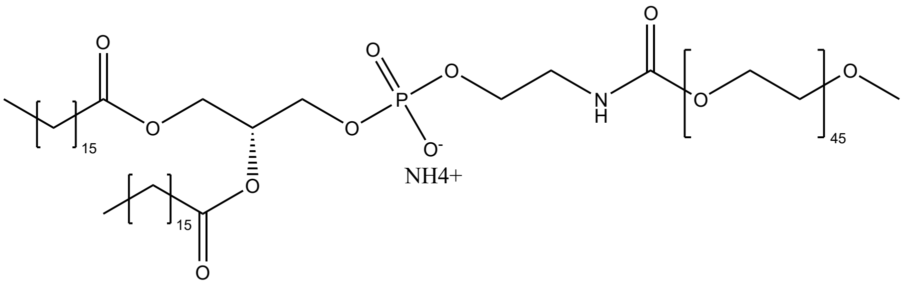 |
| | • DSPE-PEG-amine 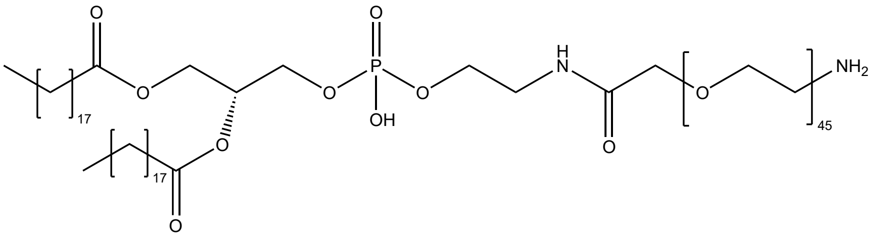 |

**Supplementary Figure S1. Chemical structures of passivation agents.**

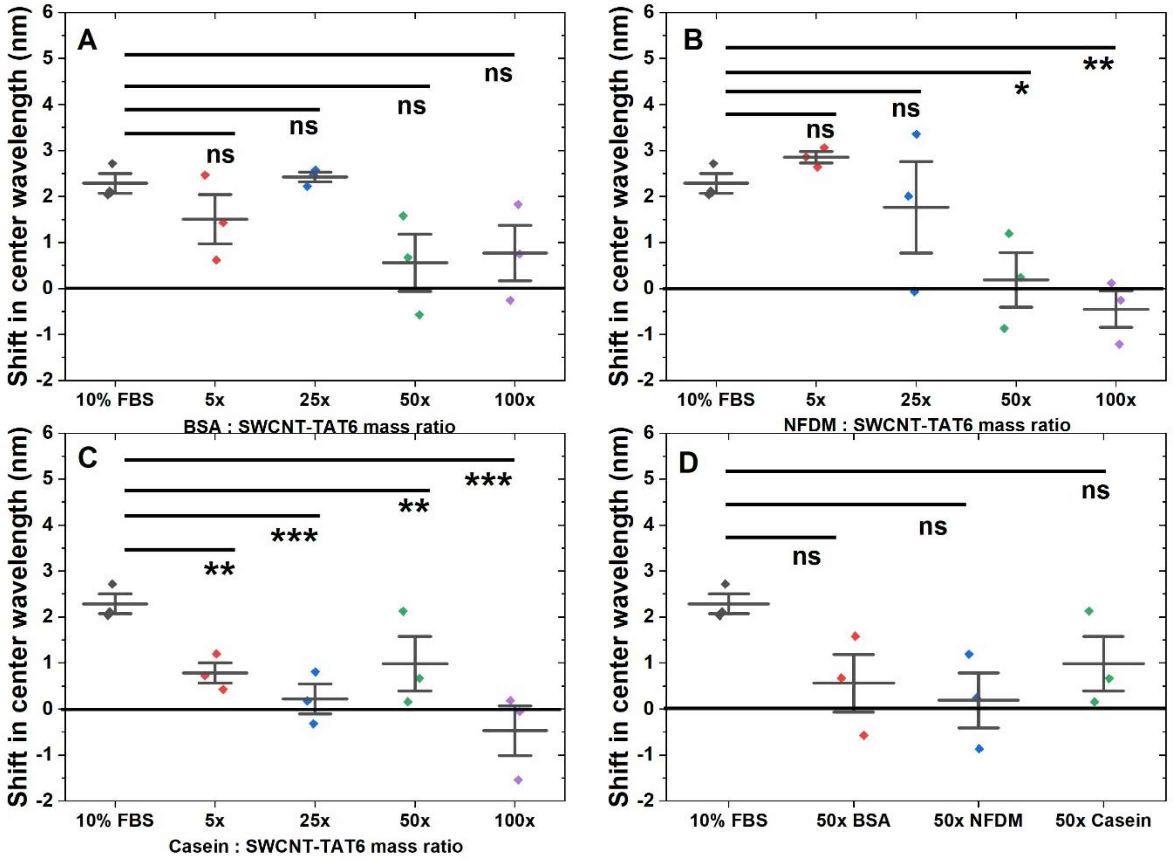


**Supplementary Figure S2. Change in (7,5) fluorescence peak upon challenging protein passivations with FBS.** (A) For BSA passivations, n=3, mean ± SD. 10% FBS (2.3 ± 0.4 nm), 5x BSA (1.5 ± 0.9 nm), 25x BSA (2.4 ± 0.2 nm), 50x BSA ( 0.6 ± 1.1 nm), 100x BSA (0.8 ± 1.0 nm).FBS and 5x BSA (0.78 nm, p=0.66), FBS and 25x BSA (0.13 nm, p=1), FBS and 50x BSA (1.7 nm, p=0.12), FBS and 100x BSA (1.5 nm, p=0.18), (B) for NFDM passivations, n=3, mean ± SD. 10% FBS (2.3 ± 0.4 nm), 5x NFDM (2.9 ± 0.2 nm), 25x NFDM (1.8 ± 1.7 nm), 50x NFDM (0.2 ± 1.0 nm), 100x NFDM (-0.4 ± 0.7 nm). FBS and 5x NFDM (0.1 nm, p=1), FBS and 25x NFDM (1.2 nm, p=0.41), FBS and 50x NFDM (2.8 nm, p=2.3E-2), FBS and 100x NFDM (3.4 nm, p=7.6E-3), (C) for casein passivations, n=3, mean ± SD. 10% FBS (2.3 ± 0.4 nm), 5x casein (0.8 ± 0.4 nm), 25x casein (0.2 ± 0.6 nm), 50x casein (1.0 ± 1.0 nm), 100x casein (-0.5 ± 0.9 nm). FBS and 5x casein (2.2 nm, p=2.66E-3), FBS and 25x casein (2.7 nm, p=6E-4), FBS and 50x casein (2 nm, p=4.8E-3), FBS and 100x casein (3.4 nm, p=1.2E-4), and (D) for 50x mass ratio, shift in emission center wavelength n=3, mean ± SD. FBS and 50x BSA (1.7 nm, p=0.2); FBS and 50x NFDM (2.1 nm, p=9.2E-2); FBS and 50x casein (1.3 nm, p=0.33).

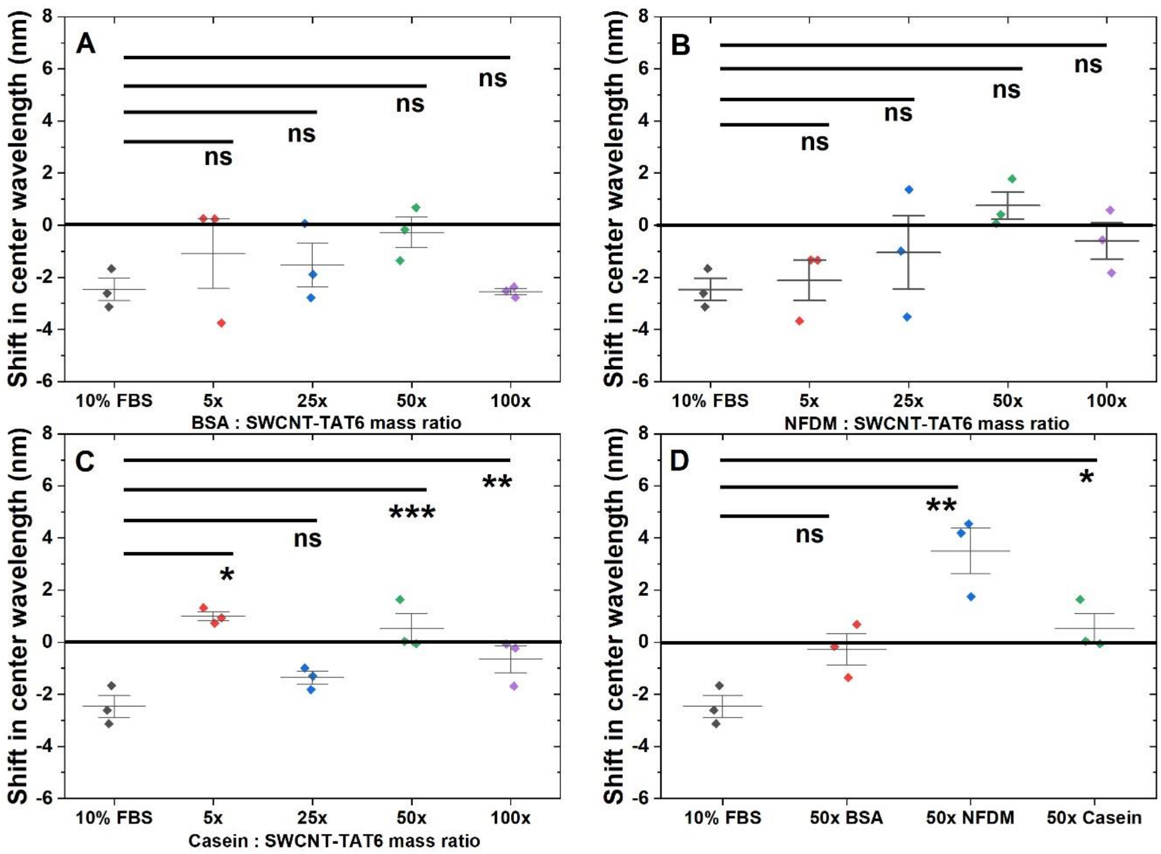


**Supplementary Figure S3. Change in (9,5) fluorescence peak upon challenging protein passivations with FBS.** (A) for BSA passivations, n=3, mean ± SD. 10% FBS (-2.5 ± 0.7 nm), 5x BSA (-1.1 ± 2.3 nm), 25x BSA (-1.5 ± 1.5 nm), 50x BSA (-0.3 ± 1 nm), 100x BSA (-2.6 ± 0.2 nm). FBS and 5x BSA (1.4 nm, p=0.54), FBS and 25x BSA (0.9 nm, p=0.8), FBS and 50x BSA (2.2 nm, p=0.2), FBS and 100x BSA (0.08 nm, p=1) (B) for NFDM passivations, n=3, mean ± SD. 10% FBS (-2.5 ± 0.7 nm), 5x NFDM (-2.1 ± 1.3 nm), 25x NFDM (-1.0 ± 2.4 nm), 50x NFDM (0.8 ± 0.9 nm), 100x NFDM (-0.6 ± 1.2 nm). FBS and 5x NFDM (0.4 nm, p=1), FBS and 25x NFDM (1.4 nm, p=0.65), FBS and 50x NFDM (3.2 nm, p=0.1), FBS and 100x NFDM (1.9 nm, p=0.4) (C) for Casein passivations, n=3, mean ± SD. 10% FBS (-2.5 ± 0.7 nm), 5x casein (1.0 ± 0.3 nm), 25x casein (-1.4 ± 0.4 nm), 50x casein (0.5 ± 1.0 nm), 100x casein (-0.7 ± 0.9 nm). FBS and 5x casein (3.8 nm, p=2E-4), FBS and 25x casein (1.4 nm, p=5.8E-2), FBS and 50x casein (3.3 nm, p=4.8E-4), FBS and 100x casein (2.1 nm, p=8E-3) (D) For 50x mass ratio, shift in emission center wavelength n=3, mean ± SD. FBS and 50x BSA (2.2 nm, p=0.1); FBS and 50x NFDM (6 nm, p=1.5E-3); FBS and 50x casein (3 nm, p=4E-2).

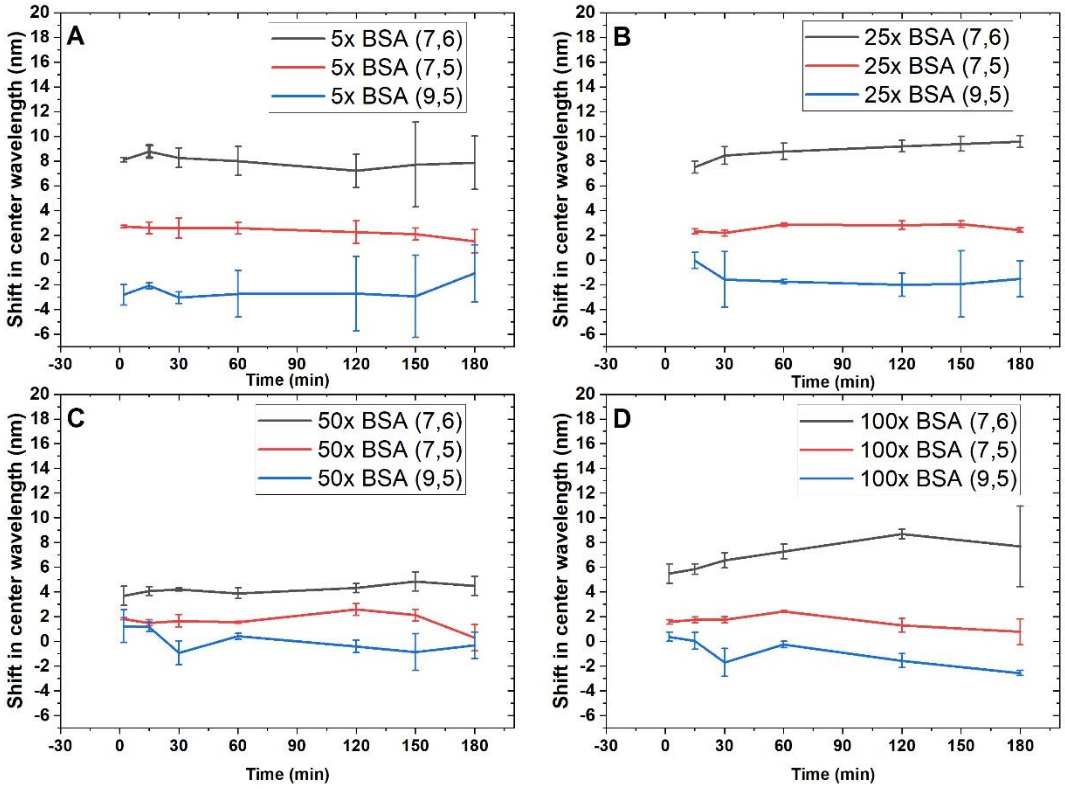


**Supplementary Figure S4. Change in all fluorescence peaks over time after addition of FBS to BSA-passivated SWCNT.** (A) For 5x BSA passivation ratio, (B) For 25x BSA passivation ratio, (C) For 50x BSA passivation ratio, (D) For 100x BSA passivation ratio.

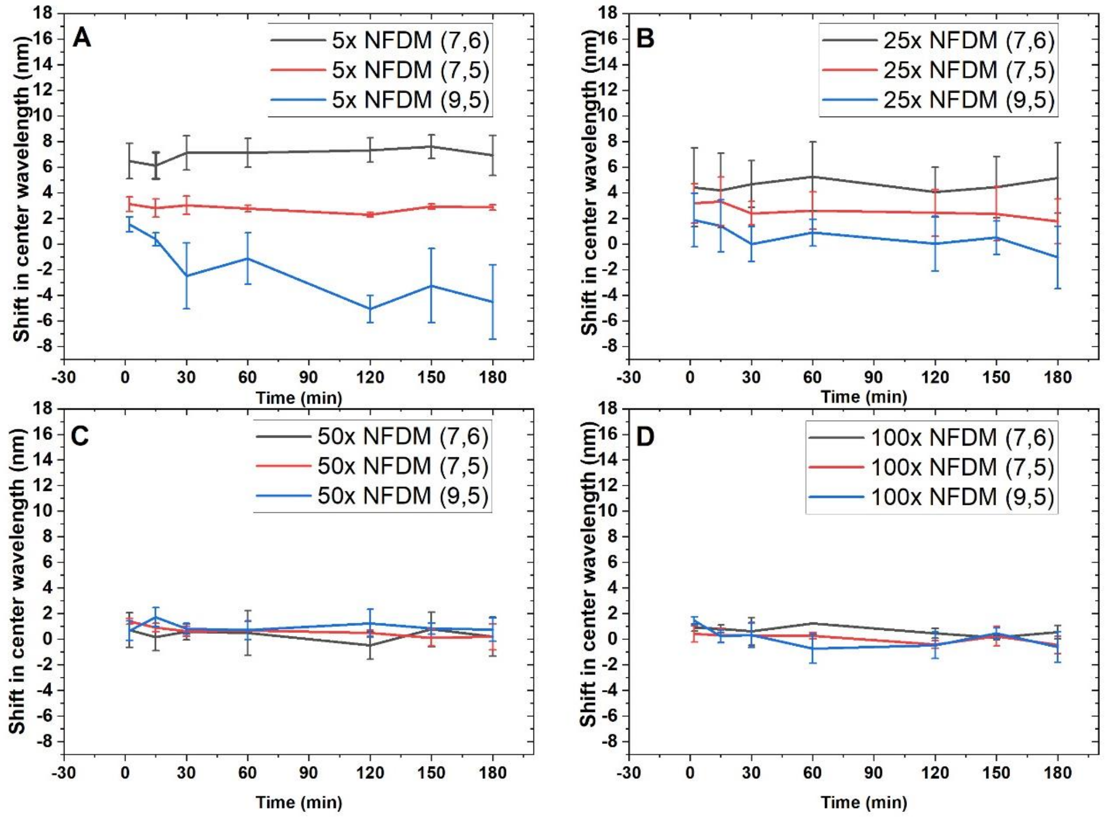


**Supplementary Figure S5. Change in all fluorescence peaks over time after addition of FBS to NFDM-passivated SWCNT.** (A) For 5x NFDM passivation ratio, (B) For 25x NFDM passivation ratio, (C) For 50x NFDM passivation ratio, (D) For 100x NFDM passivation ratio.

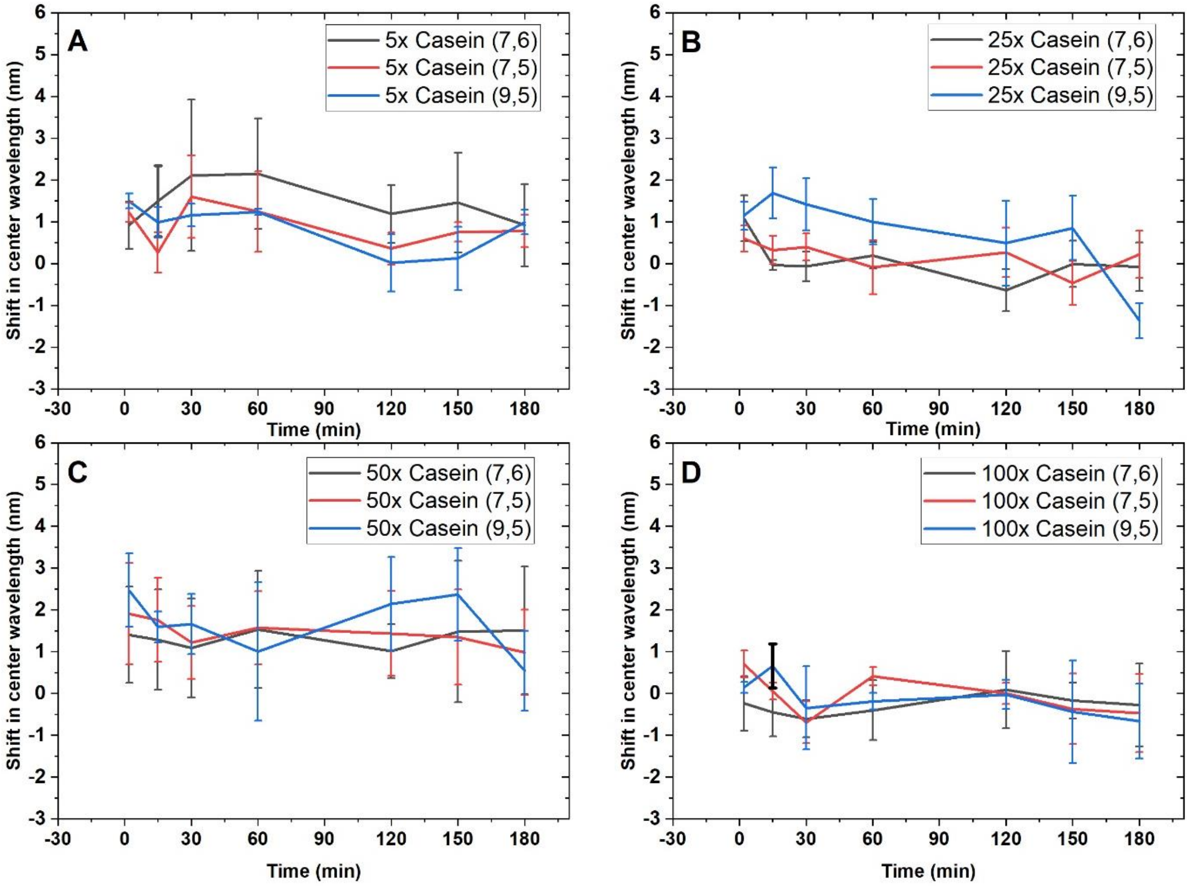


**Supplementary Figure S6. Change in all fluorescence peaks over time after addition of FBS to casein-passivated SWCNT.** (A) For 5x casein passivation ratio, (B) For 25x casein passivation ratio, (C) For 50x casein passivation ratio, (D) For 100x casein passivation ratio.

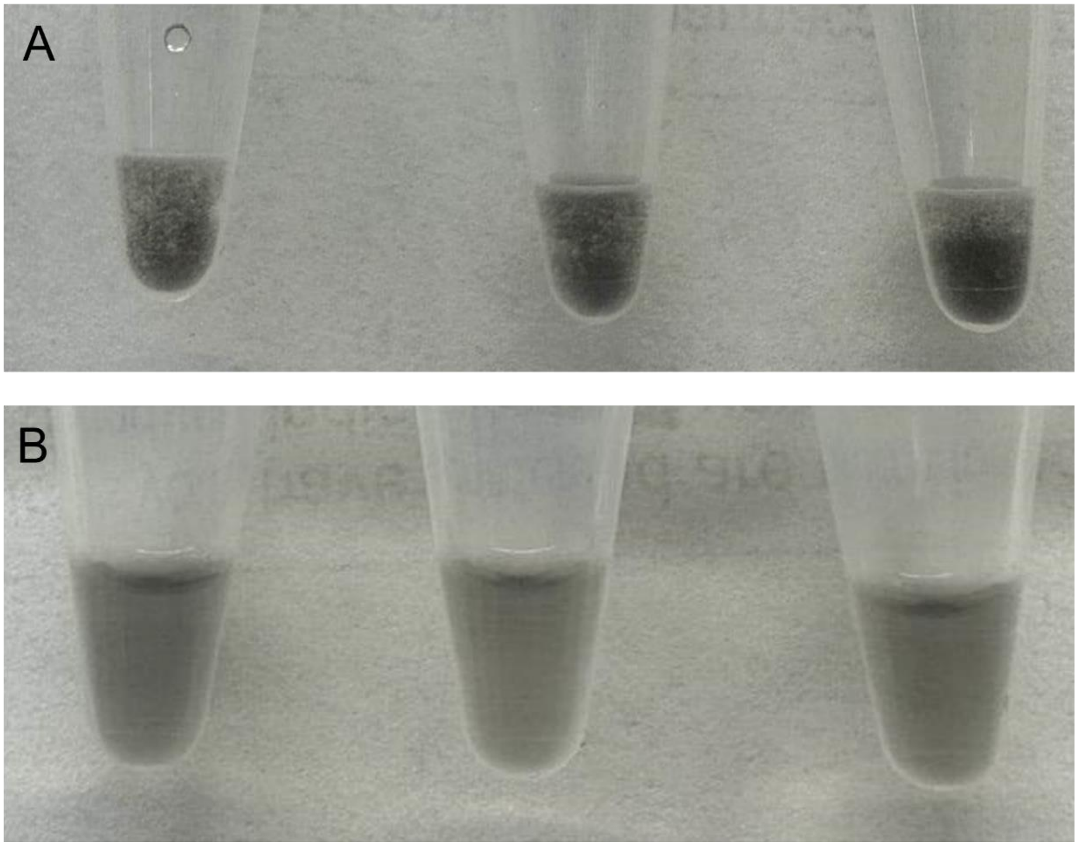


**Supplementary Figure 7. Images of SWCNT flocculation after PEI passivation.** (A) 12 hours after passivation with 50x PEI, aggregation of 10 mg/L SWCNT-$(TAT)_6$ was observed. (B) Stable solution of 50x BSA passivated 10 mg/L SWCNT-$(TAT)_6$ for at least 48 hours.

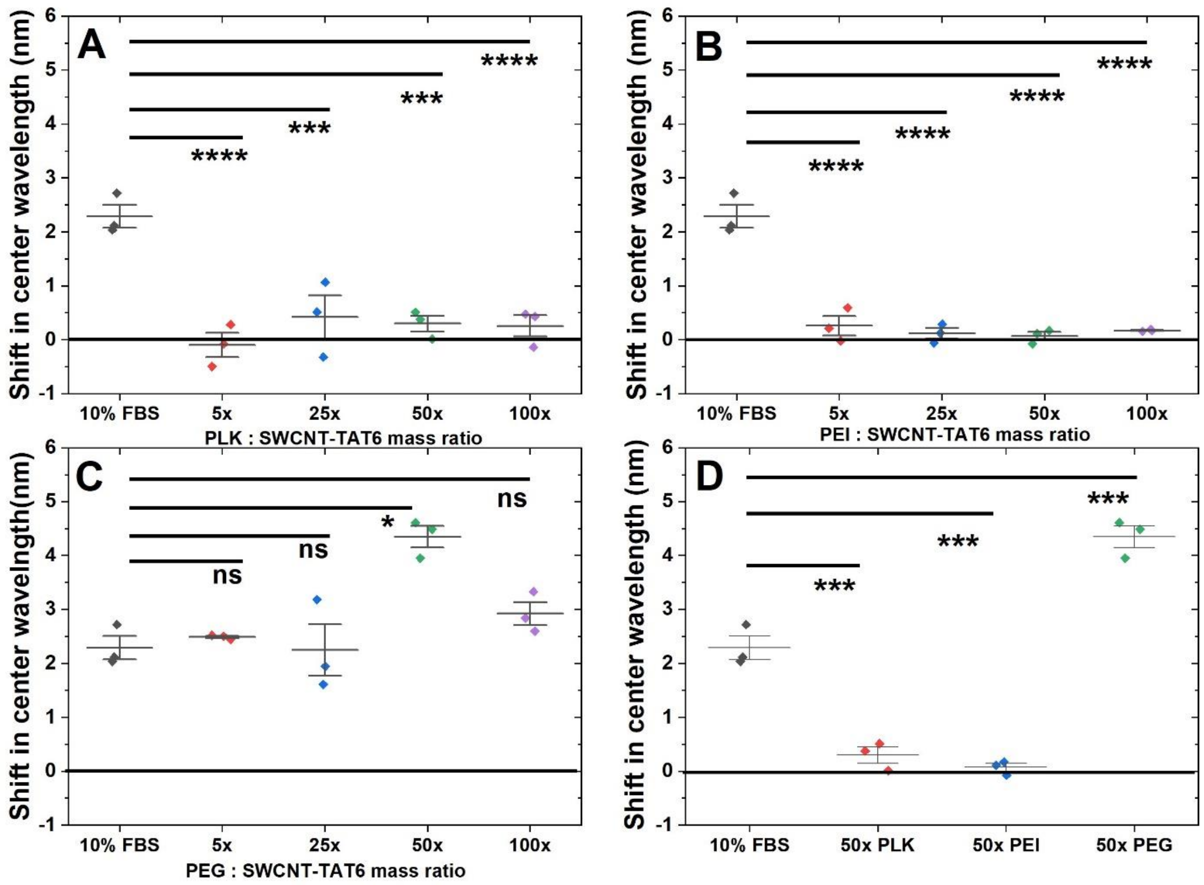


**Supplementary Figure S8. Change in (7,5) fluorescence peak upon challenging polymer passivation with FBS.** (A) For PLK passivations, n=3, mean ± SD. 10% FBS (2.3 ± 0.4 nm), 5xPLK (-0.1 ± 0.4 nm), 25x PLK (0.4 ± 0.7 nm), 50x PLK (0.3 ± 0.3 nm), 100x PLK (0.3 ± 0.3 nm). FBS and 5x PLK (3.1 nm, p=3E-5), FBS and 25x PLK (2.5 nm, p=1.3E-4), FBS and 50x PLK (2.7 nm, p=1.1E-4), FBS and 100x PLK (2.7 nm, p=9.5E-5). (B) For PEI passivations, n=3, mean ± SD. 10% FBS (2.3 ± 0.4 nm), 5x PEI (0.3 ± 0.3 nm), 25x PEI (0.1 ± 0.2 nm), 50x PEI (0.1 ± 0.1 nm), 100x PEI (0.2 ± 0.01 nm). FBS and 5x PEI (2 nm, p=2.7E-5), FBS and 25x PEI (2.2 nm, p=2.3E-5), FBS and 50x PEI (2.2 nm, p=1.9E-5), FBS and 100x PEI (2.1 nm, p=1.8E-5). (C) For PEG passivations, n=3, mean ± SD. 10% FBS (2.3 ± 0.4 nm), 5x PEG (2.5 ± 0.04 nm), 25x PEG (2.2 ± 0.8 nm), 50x PEG (4.3 ± 0.3 nm), 100x PEG (2.9 ± 0.4 nm). FBS and 5x PEG (0.5 nm, p=0.6), FBS and 25x PEG (0.7 nm, p=0.28), FBS and 50x PEG (1.4 nm, p=0.03), FBS and 100x PEG (0.04 nm,p=1). (D) For 50x mass ratio, shift in emission center wavelength n=3, mean ± SD. FBS and 50x PLK (2 nm, p=7.8E-4); FBS and 50x PEI (2.2 nm, p=4E-4); FBS and 50x PEG (2.1 nm, p=6.3E-4).

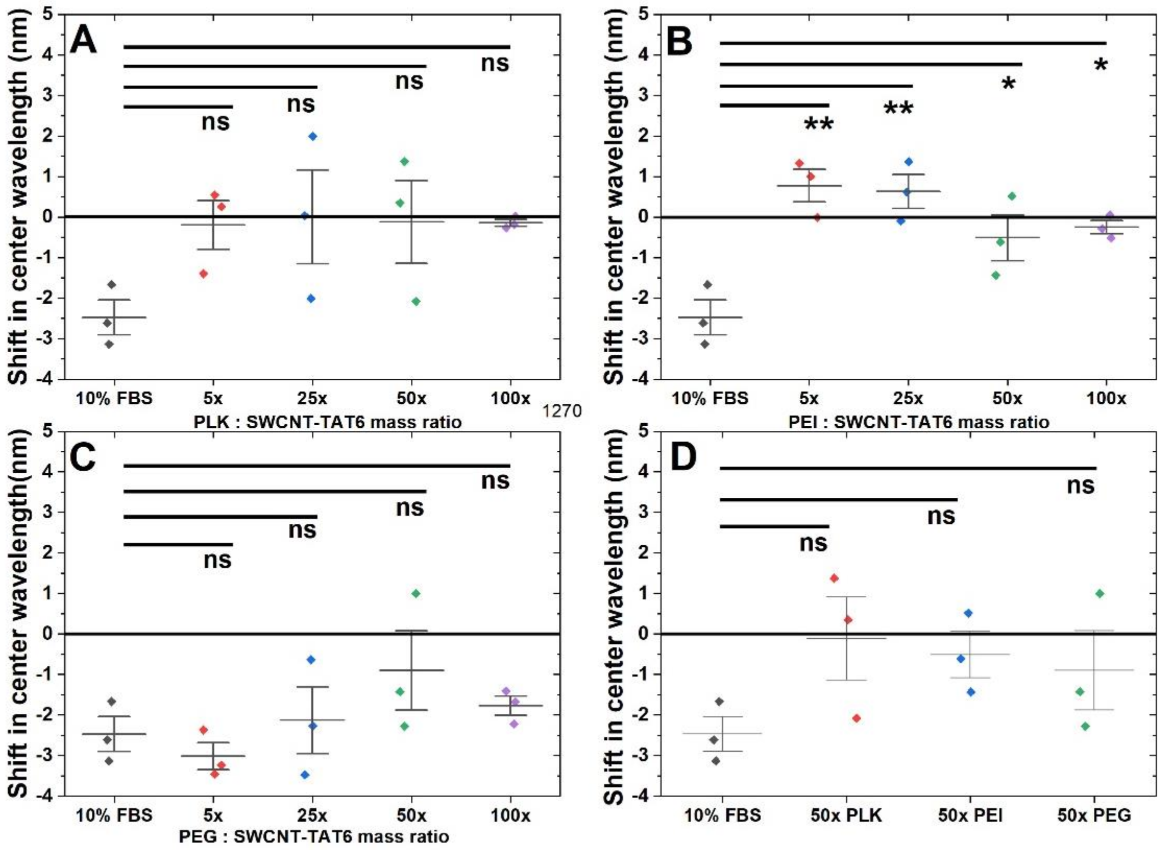


**Supplementary Figure S9. Change in (9,5) fluorescence peak upon challenging polymer passivation with FBS.** (A) For PLK passivations, n=3, mean ± SD. 10% FBS (-2.5 ± 0.7 nm), 5x PLK (-0.2 ± 1.0 nm), 25x PLK (0.01 ± 2.0 nm), 50x PLK (-0.1 ± 1.8 nm), 100x PLK (-0.1 ± 0.1 nm). FBS and 5x PLK (2.6 nm, p=0.1), FBS and 25x PLK (2.8 nm, p=7.6E-2), FBS and 50x PLK (2.7 nm, p=0.1), FBS and 100x PLK (2.7 nm, p=0.1). (B) For PEI passivations, n=3, mean ± SD. 10% FBS (-2.5 ± 0.7 nm), 5x PEI (0.8 ± 0.7 nm), 25x PEI (0.6 ± 0.7 nm), 50x PEI (-0.5 ± 1.0 nm), 100x PEI (-0.2 ± 0.3 nm). FBS and 5x PEI (3.6 nm, p=1.3E-3), FBS and 25x PEI (3.4 nm, p =1.7E-3), FBS and 50x PEI 2.3 nm, p=1.8E-2), FBS and 100x PEI (2.6 nm, p=1E-2). (C) For PEG passivations, n=3, mean ± SD. 10% FBS (-2.5 ± 0.7 nm), 5x PEG (-3.0 ± 0.6 nm), 25x PEG (-2.1 ± 1.4 nm), 50x PEG (-0.9 ± 1.7 nm), 100x PEG (-1.8 ± 0.4 nm). FBS and 5x PEG (0.2 nm, p=1), FBS and 25x PEG (0.7 nm, p=0.85), FBS and 50x PEG (1.9 nm, p=0.2), FBS and 100x PEG (1.04 nm, p=0.61). (D) For 50x mass ratio, shift in emission center wavelength n=3, mean ± SD. FBS and 50x PLK (2.7 nm, p=0.16); FBS and 50x PEI (2.3 nm, p=0.24); FBS and 50x PEG (1.9 nm, p=0.4).

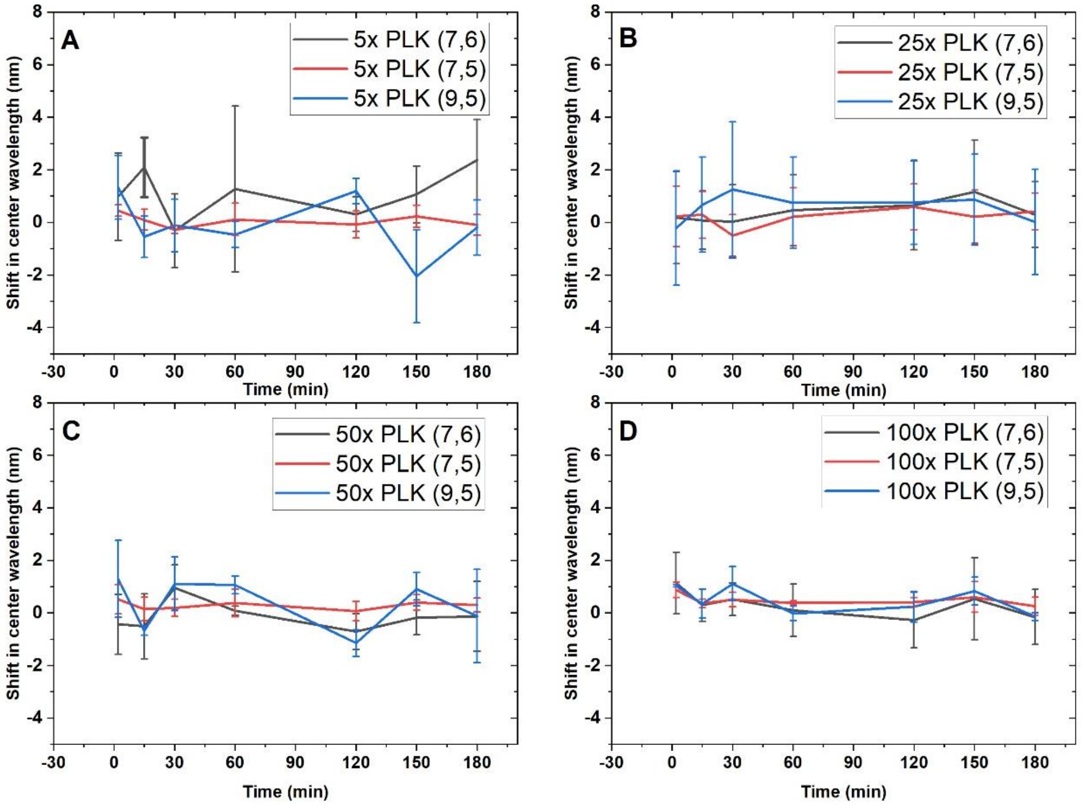


**Supplementary Figure S10. Change in all fluorescence peaks over time after passivation after FBS addition for PLK-passivated SWCNT** (A) For 5x PLK passivation ratio, (B) For 25x PLK passivation ratio, (C) For 50x PLK passivation ratio, (D) For 100x PLK passivation ratio.

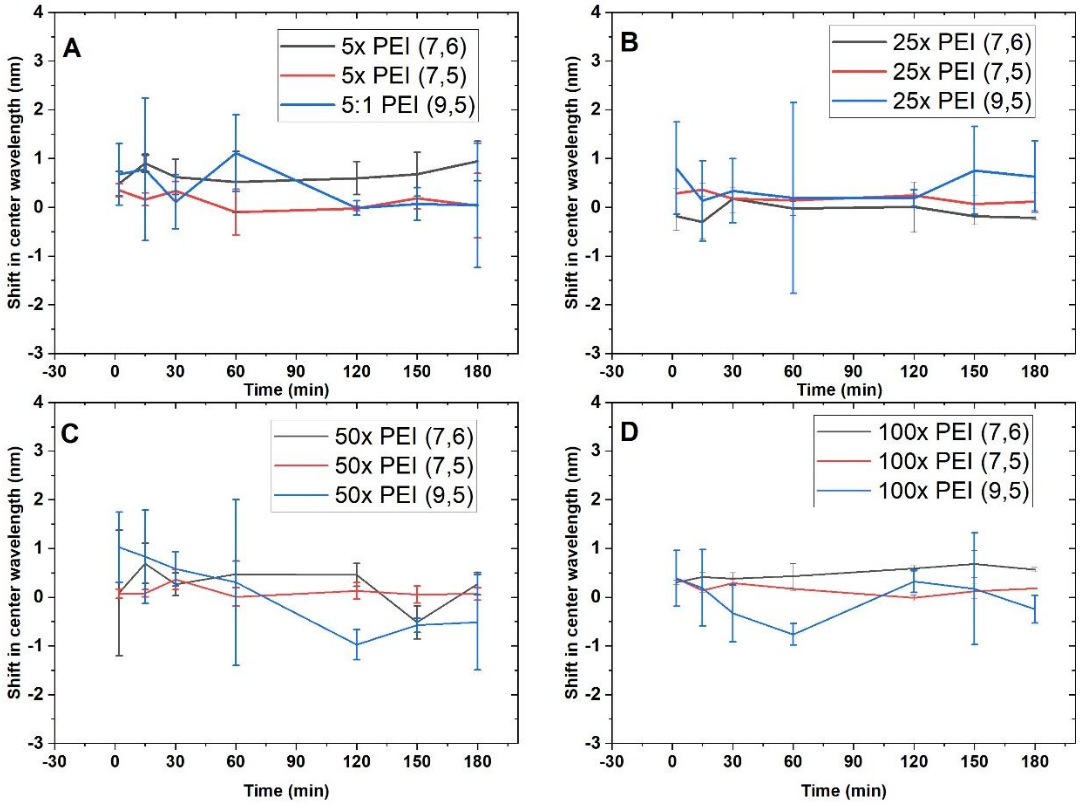


**Supplementary Figure S11. Change in all fluorescence peaks over time after passivation after FBS addition for PEI-passivated SWCNT.** (A) For 5x PEI passivation ratio, (B) For 25x PEI passivation ratio, (C) For 50x PEI passivation ratio, (D) For 100x PEI passivation ratio

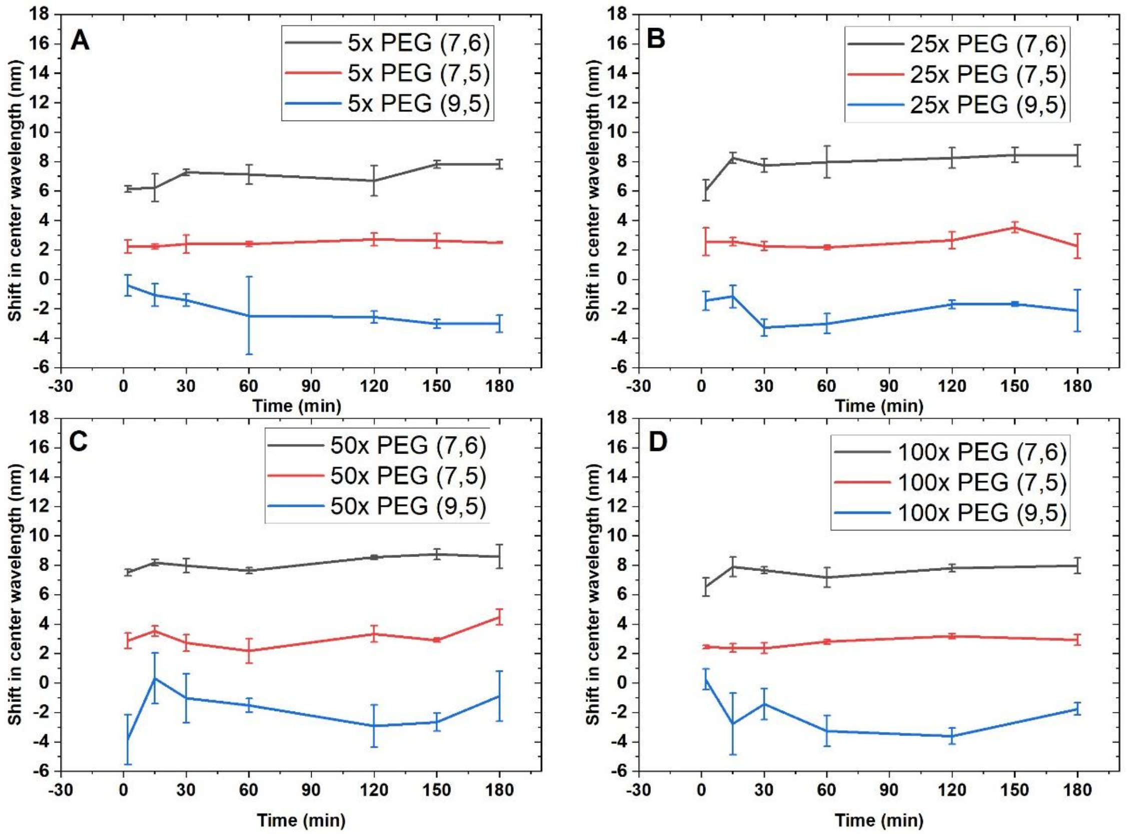


**Supplementary Figure S12. Change in all fluorescence peaks over time after addition of FBS for PEG-passivated SWCNT.** (A) For 5x PEG passivation ratio, (B) For 25x PEG passivation ratio, (C) For 50x PEG passivation ratio, (D) For 100x PEG passivation ratio.

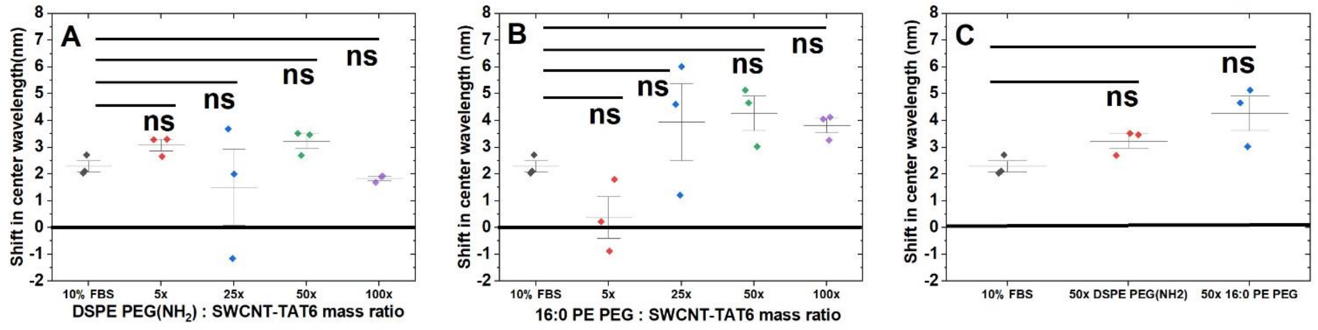


**Supplementary Figure S13. Change in (7,5) fluorescence peak upon challenging phospholipid passivation with FBS.** (A) For DSPE PEG ($NH_2$) passivations, n=3, mean ± SD. 10% FBS (2.3 ± 0.4 nm), 5x DSPE PEG ($NH_2$) (3.1 ± 0.4 nm), 25x DSPE PEG ($NH_2$) (1.5 ± 2.5 nm), 50x DSPE PEG ($NH_2$) (3.2 ± 0.5 nm), 100x DSPE PEG ($NH_2$) (1.8 ± 0.1 nm). FBS and 5x DSPE PEG ($NH_2$) (0.8 nm, p=0.8), FBS and 25x DSPE PEG ($NH_2$) (0.8, p=0.82), FBS and 50x DSPE PEG ($NH_2$) (0.95 nm, p=0.7), FBS and 100x DSPE PEG ($NH_2$) (0.4 nm, p=1). (B) For 16:0 PE PEG passivations, n=3, mean ± SD. 10% FBS (2.3 ± 0.4 nm), 5x 16:0 PE PEG (0.4 ± 1.3 nm), 25x 16:0 PE PEG (3.9 ± 2.5 nm), 50x 16:0 PE PEG (4.3 ± 1.1 nm), 100x 16:0 PE PEG (3.8 ± 0.5 nm). FBS and 5x 16:0 PE PEG (2.6 nm, p=0.2), FBS and 25x 16:0 PE PEG (0.98 nm, p=0.8), FBS and 50x 16:0 PE PEG (1.3 nm, p=0.7), FBS and 100x 16:0 PE PEG (0.9 nm, p=0.9). (C) For 50x mass ratio, shift in emission center wavelength n=3, mean ± SD. FBS and 50x DSPE PEG ($NH_2$) (0.3, p=0.9); FBS and 50x 16:0 PE PEG (1.3 nm, p=0.14).

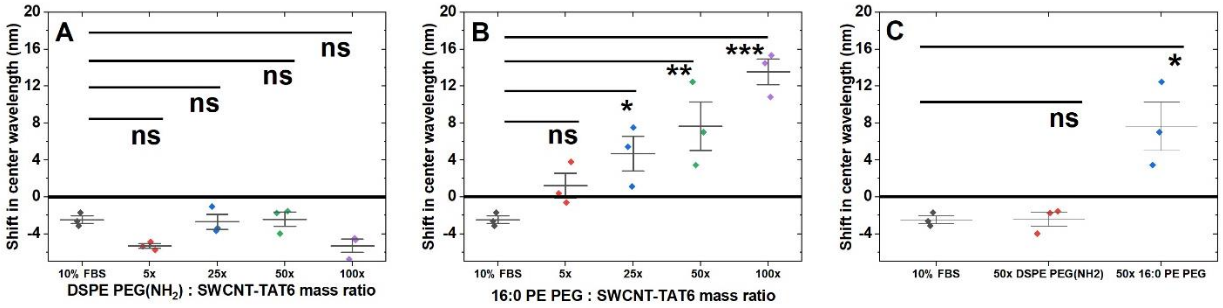


**Supplementary Figure S14 Change in (9,5) fluorescence peak upon challenging phospholipid passivation with FBS.** (A) For DSPE PEG ($NH_2$) passivations, n=3, mean ± SD. 10% FBS (-2.5 ± 0.7 nm), 5x DSPE PEG ($NH_2$) (-5.3 ± 0.4 nm), 25x DSPE PEG ($NH_2$) (-2.7 ± 1.4 nm), 50x DSPE PEG ($NH_2$) (-2.4 ± 1.3 nm), 100x DSPE PEG ($NH_2$) (-5.3 ± 1.2 nm). FBS and 5x DSPE PEG ($NH_2$) (2.5 nm, p=0.1), FBS and 25x DSPE PEG ($NH_2$) (0.1 nm, p=1), FBS and 50x DSPE PEG ($NH_2$) (0.4 nm, p=0.98), FBS and 100x DSPE PEG ($NH_2$) (2.5 nm, p=0.1). (B) For 16:0 PE PEG passivations, n=3, mean ± SD. 10% FBS (-2.5 ± 0.7 nm), 5x 16:0 PE PEG (1.2 ± 2.3 nm), 25x 16:0 PE PEG (4.7 ± 3.2 nm), 50x 16:0 PE PEG (7.7 ± 4.5 nm), 100x 16:0 PE PEG (13.6 ± 2.4 nm). FBS and 5x 16:0 PE PEG (4 nm, p=0.4), FBS and 25x 16:0 PE PEG (7.5 nm, p=4.7E-2), FBS and 50x 16:0 PE PEG (10.5 nm, p=8.9E-3), FBS and 100x 16:0 PE PEG (16.4 nm, p=5.4E-4). (C) For 50x mass ratio, shift in emission center wavelength n=3, mean ± SD. FBS and 50x DSPE PEG ($NH_2$) (0.4, p=0.9); FBS and 50x 16:0 PE PEG (10.5 nm, p=1.4E-2).

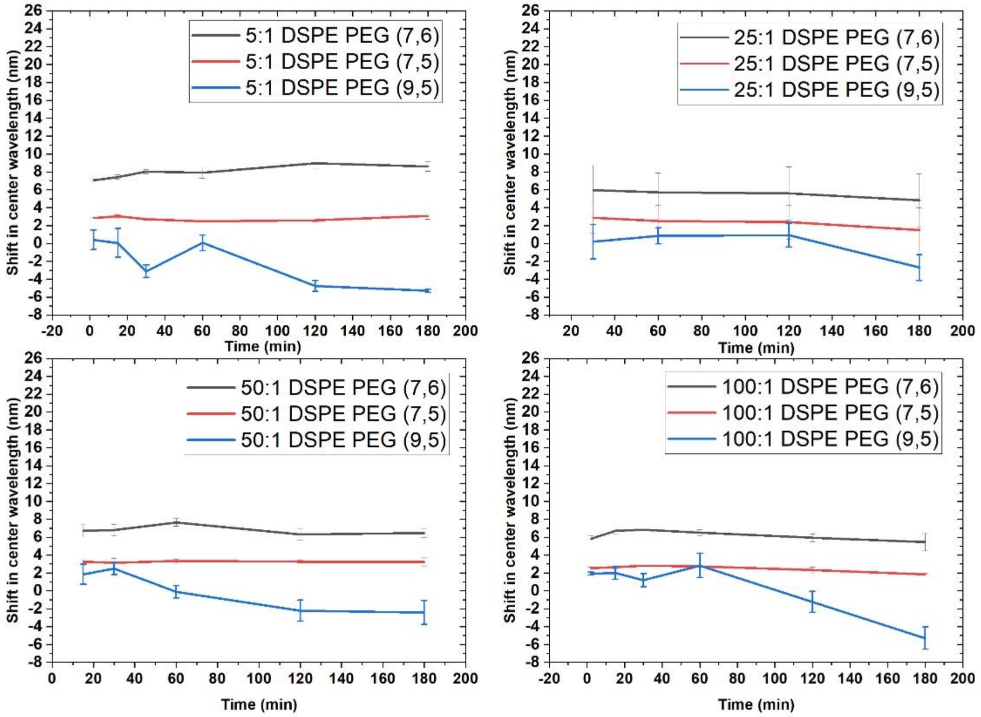


**Supplementary Figure S15. Change in all fluorescence peaks over time after addition of FBS to DSPE-PEG ($NH_2$)-passivated SWCNT.** (A) For 5x DSPE PEG ($NH_2$) passivation ratio, (B) For 25x DSPE PEG ($NH_2$) passivation ratio, (C) For DSPE PEG ($NH_2$) 50x passivation ratio, (D) For DSPE PEG ($NH_2$) 100x passivation ratio.

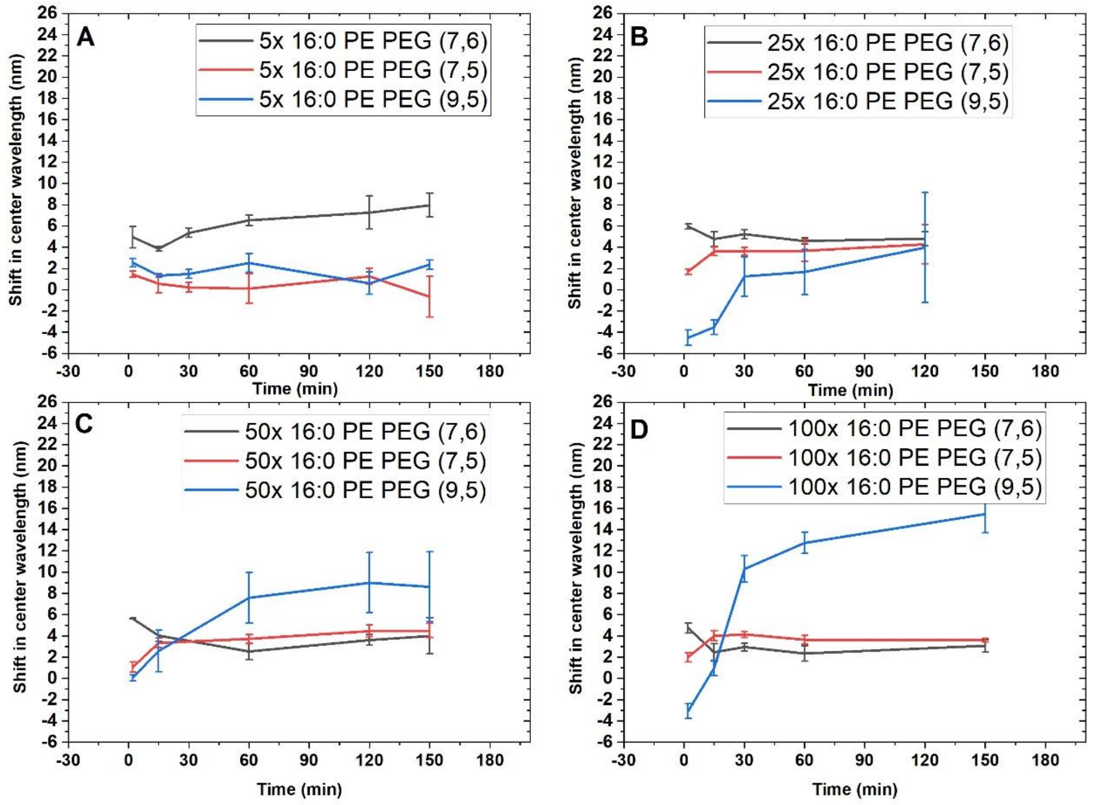


**Supplementary Figure S16. Change in all fluorescence peaks over time after FBS addition to PE2000PEG-passivated SWCNT.** (A) For 5x PE2000PEG passivation ratio. (B) For 25x DSPE PE2000PEG passivation ratio, (C) For 50x PE2000PEG passivation ratio, and (D) For 100x DSPE PEG PE2000PEG passivation ratio.

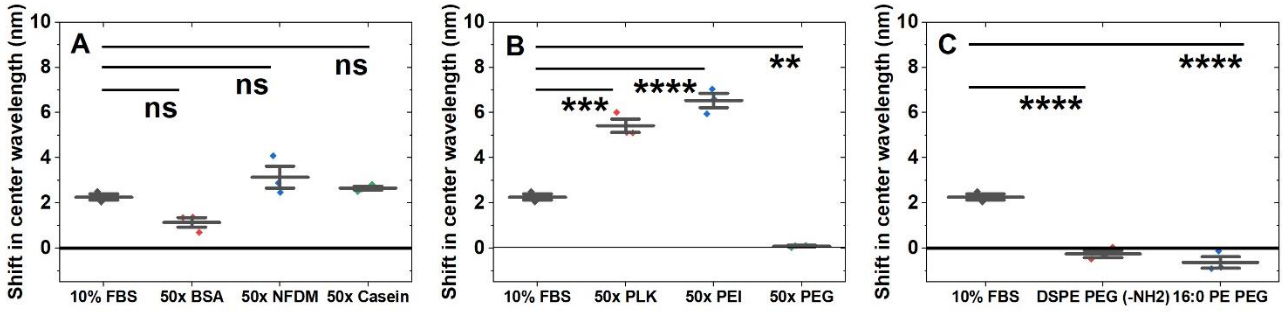


**Supplementary Figure S17. Change in (7,5) absorbance peak of SWCNT-$(TAT)_6$ in presence of passivation agents compared to FBS alone.** (A) For protein passivations, n=3, mean ± SD. (10% FBS = 2.3 ± 0.2 nm), 50x BSA (1.1 ± 0.4 nm), 50x NFDM (3.2 ± 0.8 nm), 50x casein (2.7 ± 0.2 nm). FBS and 50x BSA (1.1 nm, p=0.08), FBS and 50x NFDM (0.9 nm, p=0.2), FBS and 50x casein (0.4 nm, p=0.7). (B) For polymer passivations, n=3, mean ± SD. (10% FBS = 2.3 ± 0.2 nm), 50x PLK (5.4 ± 0.5 nm), 50x PEI (7.3 ± 0.9 nm), 50x PEG (0.1 ± 0.1 nm). FBS and 50x PLK (3.1 nm, p=2.1E-4), FBS and 50x PEI (4.3 nm, p=9E-6), FBS and 50x PEG (2.2 nm, p=2.2E-3). (C) For phospholipid passivations, n=3, mean ± SD. (10% FBS = 2.3 ± 0.2 nm), 50x DSPE PEG ($NH_2$) (-0.3 ± 0.3 nm) and 50x 16:0 PE PEG (-0.6 ± 0.4). FBS and 50x DSPE PEG ($NH_2$) (2.5 nm, p=2.8E-5), and FBS and 50x 16:0 PE PEG (2.9 nm, p=1.6E-5).

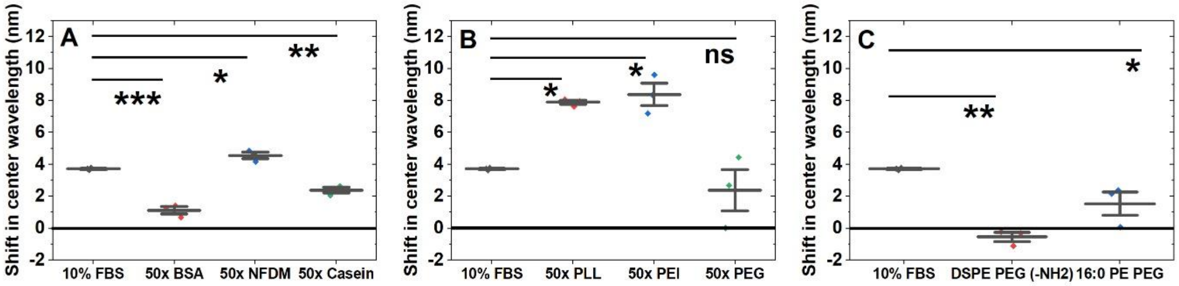


**Supplementary Figure S18. Change in (9,5) absorbance peak of SWCNT-$(TAT)_6$ in the presence of passivation agents compared to FBS.** (A) For protein passivations, n=3, mean ± SD. (10% FBS = 3.7 ± 0.1 nm), 50x BSA (1.1 ± 0.4 nm), 50x NFDM (4.56 ± 0.35 nm), 50x casein (2.4 ± 0.3 nm). FBS and 50x BSA (2.6 nm, p=1.4E-4), FBS and 50x NFDM (0.8 nm, p=6.1E-2), FBS and 50x casein (1.3 nm, p=8.5E-3). (B) For polymer passivations, n=3, mean ± SD. (10% FBS = 3.7 ± 0.1 nm), 50x PLK (7.9 ± 0.2 nm), 50x PEI (8.9 ± 2.1 nm), 50x PEG (-2.4 ± 2.2 nm). FBS and 50x PLK (4.2 nm, p=2.3E-2), FBS and 50x PEI (4.7 nm, p=1.4E-2), FBS and 50x PEG (1.34 nm, p=0.53). (C) For phospholipid passivations, n=3, mean ± SD. (10% FBS = 3.7 ± 0.1 nm), 50x DSPE PEG ($NH_2$) (-5.5 ± 0.5 nm) and 50x 16:0 PE PEG (1.5 ± 1.3 nm). FBS and 50x DSPE PEG ($NH_2$) (4.3 nm, p=1.8E-3), and FBS and 50x 16:0 PE PEG (2.2 nm, p=2.1E-2).

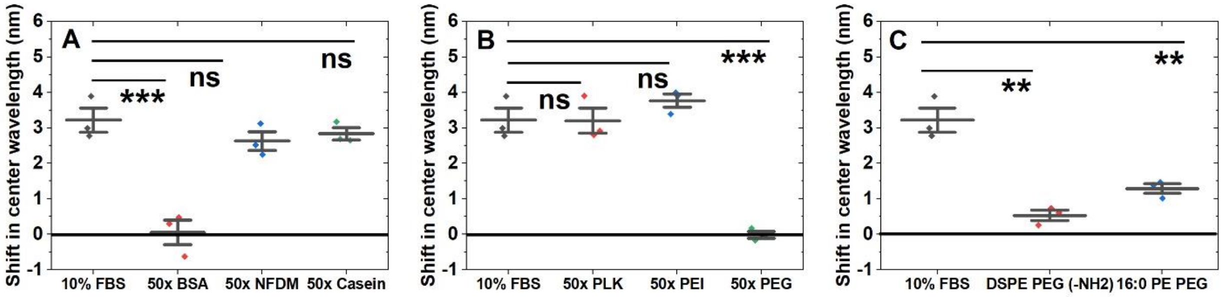


**Supplementary Figure S19. Change in (6,5) absorbance peak of SWCNT-$(TAT)_6$ in the presence of passivation agents compared to FBS.** (A) For protein passivations, n=3, mean ± SD. (10% FBS = 3.2 ± 0.6 nm), 50x BSA (0.05 ± 0.6 nm), 50x NFDM (2.6 ± 0.4 nm), 50x casein (2.8 ± 0.3 nm). FBS and 50x BSA (3.2 nm, p=9.5E-4), FBS and 50x NFDM (0.6 nm, p=0.46), FBS and 50x casein (0.38 nm, p=0.7). (B) For polymer passivations, n=3, mean ± SD. (10% FBS = 3.22 ± 0.6 nm), 50x PLK (3.2 ± 0.6 nm), 50x PEI (5.2 ± 1 nm), 50x PEG (-1.2 ± 0.2 nm). FBS and 50x PLK (1.5E-2 nm, p=1), FBS and 50x PEI (0.55 nm, p=0.44), FBS and 50x PEG (3.2 nm, p=4.4E-4). (C) For phospholipid passivations, n=3, mean ± SD. (10% FBS = 3.2 ± 0.6 nm), 50x DSPE PEG ($NH_2$) (5.4 ± 0.3 nm) and 50x 16:0 PE PEG (1.3 ± 0.2 nm). FBS and 50x DSPE PEG ($NH_2$) (2.7 nm, p=3E-3), and FBS and 50x 16:0 PE PEG (1.9 nm, p=1E-2).

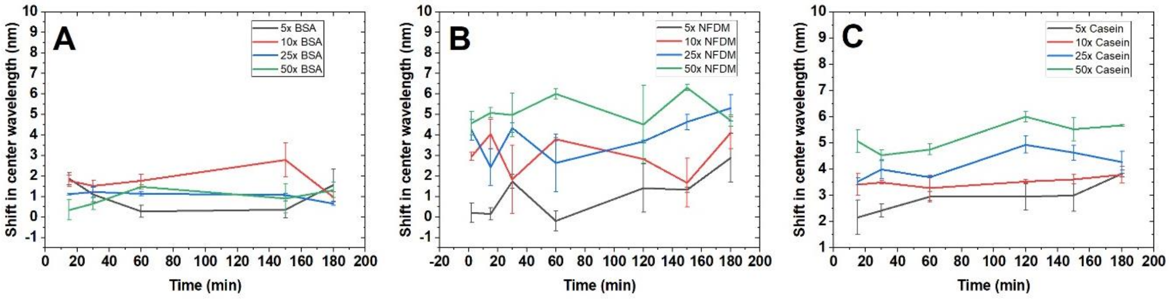


**Supplementary Figure S20. Change in center wavelength of (7,6) chirality absorption peak over time after addition of protein passivation agents.** (A) For all BSA mass ratio passivation, (B) For all Non-fat dry milk mass ratio passivations, and (C) For all casein mass ratio passivations.

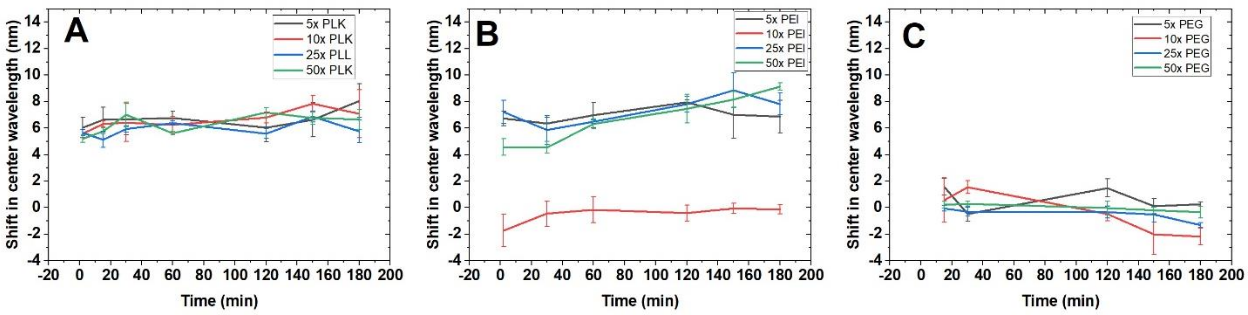


**Supplementary Figure S21. Change in center wavelength of (7,6) chirality absorption peak over time after addition of polymer passivation agents.** (A) For all poly-L-lysine mass ratio passivations, (B) For all polyethylene imine mass ratio passivations, and (C) For all polyethylene glycol mass ratio passivations.

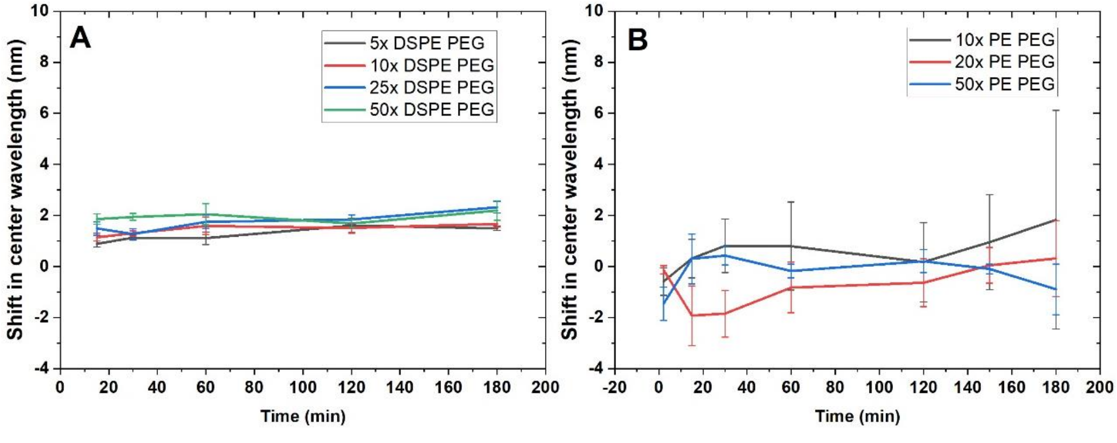


**Supplementary Figure S22. Change in center wavelength of (7,6) chirality absorption peak over time after addition of phospholipid passivation agents.** (A) For all DSPE PEG ($NH_2$) mass ratio passivations and (B) For all 16:0 PE 2000 PEG mass ratio passivations.

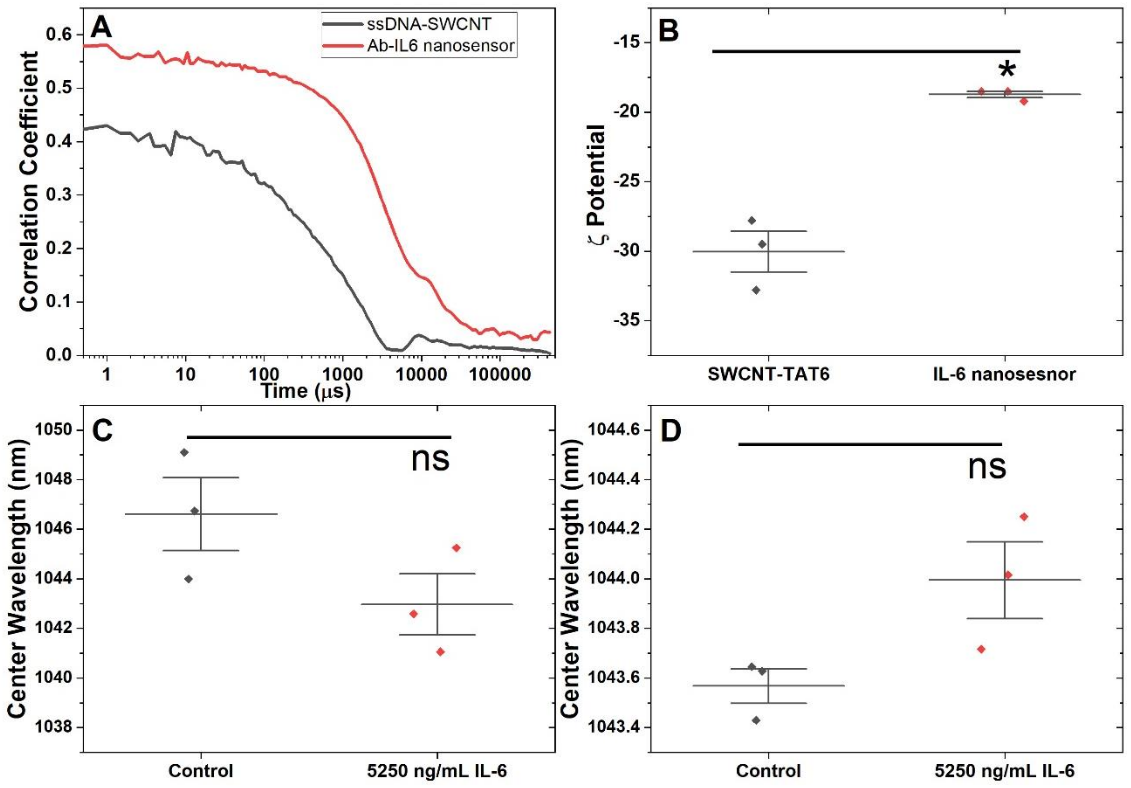


**Supplementary Figure S23. Characterization and in vitro performance of an engineered IL-6 nanosensor.** Successful conjugation of the antibody-based IL-6 nanosensor was assessed by (A) comparison of decay in correlation coefficient as function of time for SWCNT-$(TAT)_6$ and IL-6 Antibody (Ab) conjugated SWCNT-$(TAT)_6$, (B) Change in zeta potential for SWCNT-$(TAT)_6$ (-30 ± 2.5 mV) compared to the IL-6 nanosensor (-18.7 ± 0.4 mV) (difference in means = -11.3 mV, p=0.01, two-tailed t-test). (C) Performance of the IL-6 nanosensor was assessed by comparing change in the emission peak for IL-6 nanosensor in presence and absence of IL-6 in 1x PBS, SWCNT-$(TAT)_6$ (1046.6 ± 2.6 nm) and Ab IL-6 conjugated (1043 ± 2.1 nm) (difference in means = 3.64 nm, p=0.13, two-tailed t-test). (D) Performance of the IL-6 nanosensor in 10% was assessed by comparing the change in the emission peak for IL-6 nanosensor in the presence and absence of IL-6 in 10% FBS, SWCNT-$(TAT)_6$ (1043.6 ± 0.1 nm) and Ab IL-6 conjugated (1044 ± 0.3 nm) (difference in means = -0.43 nm, p=0.07, two-tailed t-test).

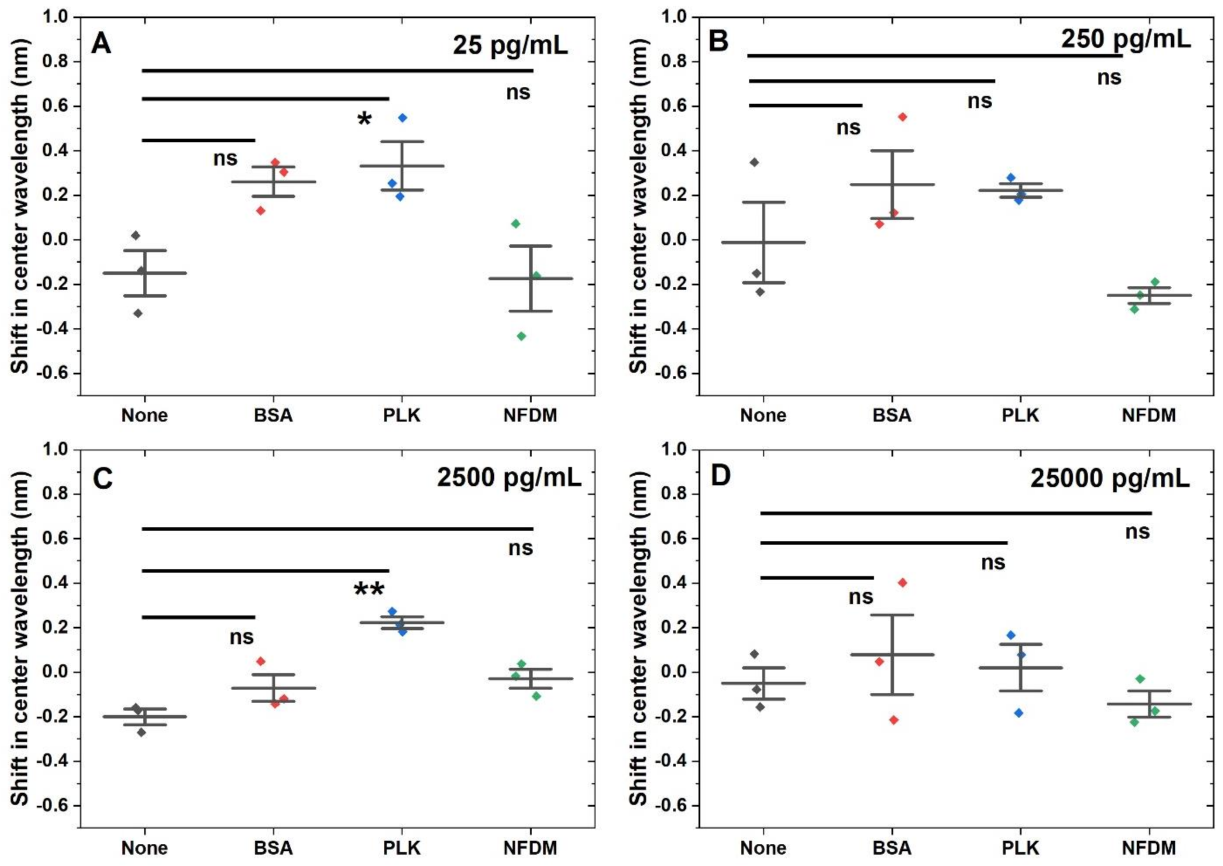


**Supplementary Figure 24. Response of the (7,5) nanosensor chirality to IL-6 in human serum.** Shift in emission center wavelength for (7,5) chirality of IL-6 nanosensor to (A) 25 pg/mL IL-6. n=3, mean ± SD. Non-passivated (-0.15 ± 0.05 nm), 50x BSA passivated (0.26 ± 0.11 nm), 50x PLK passivated (0.33 ± 0.19 nm), and 50x NFDM passivated (-0.17 ± 0.02 nm). For groups, Non-passivated and 50x BSA passivated nanosensor (0.41 nm, p=7.6E-2); Non-passivated and 50x PLK (0.48 nm, p=4.2E-2); Non-passivated and 50x NFDM (-0.02 nm, p=0.997). (B) Nanosensor response to 250 pg/mL IL-6. n=3, mean ± SD. Non-passivated (-0.01 ± 0.3 nm), 50x BSA passivated (0.25 ± 0.26 nm), 50x PLK passivated (0.22 ± 0.05 nm), and 50x NFDM passivated (-0.25 ± 0.06 nm). For groups, Non-passivated and 50x BSA passivated nanosensor (0.26 nm, p=0.24); Non-passivated and 50x PLK (0.23 nm, p=0.3); Non-passivated and 50x NFDM (-0.24 nm, p=0.3). (C) Nanosensor response to 2,500 pg/mL IL-6. n=3, mean ± SD. Non-passivated (-0.2 ± 0.06 nm), 50x BSA passivated (-0.07 ± 0.1 nm), 50x PLK passivated (0.22 ± 0.05 nm), and 50x NFDM passivated (-0.03 ± 0.07 nm). For groups, Non-passivated and 50x BSA passivated nanosensor (0.13 nm, p=0.17); Non-passivated and 50x PLK (0.4 nm, p=1.1E-3); Non-passivated and 50x NFDM (0.17 nm, p=7.2E-2). (D) Nanosensor response to 25,000 pg/mL IL-6. n=3, mean ± SD. Non-passivated (-0.05 ± 0.04 nm), 50x BSA passivated (0.08 ± 0.31 nm), 50x PLK passivated (0.02 ± 0.2 nm), and 50x NFDM passivated (-0.14 ± 0.08 nm). For groups, Non-passivated and 50x BSA passivated nanosensor (0.13 nm, p=0.8); Non-passivated and 50x PLK (0.07 nm, p=0.95); Non-passivated and 50x NFDM (-0.09 nm, p=0.91).

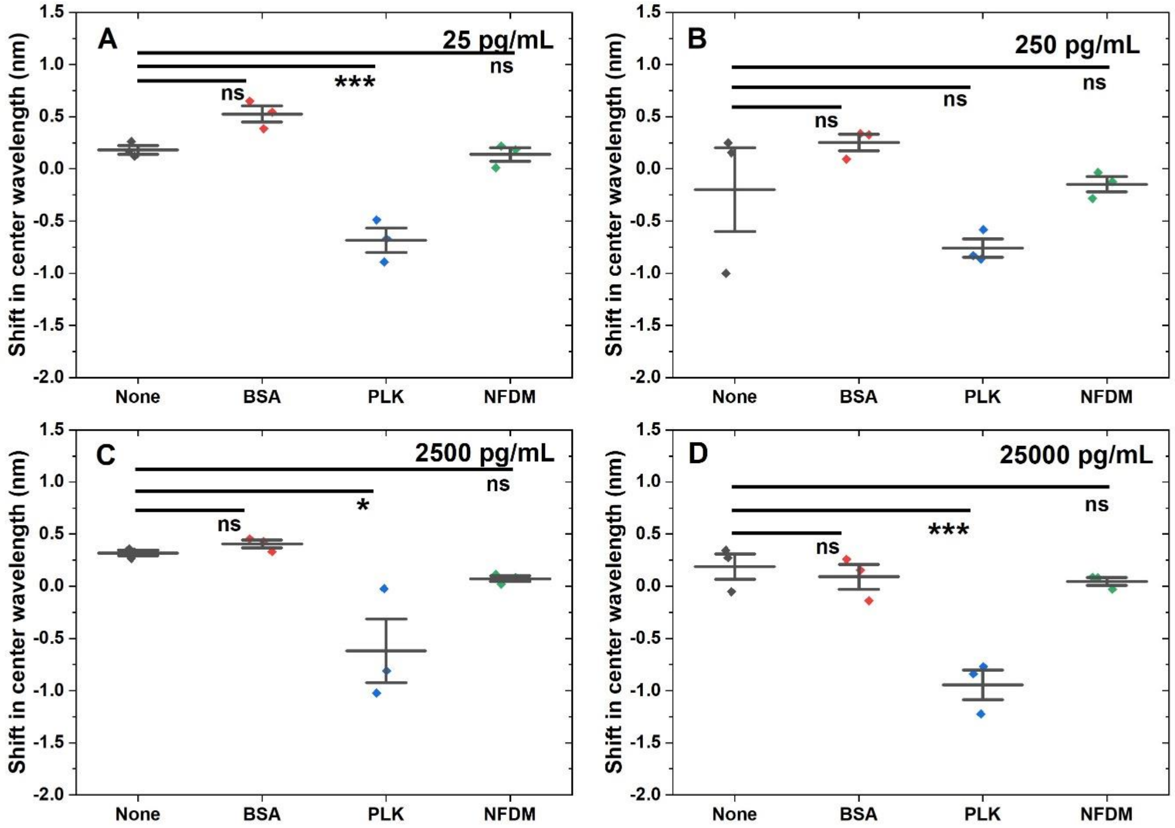


**Supplementary Figure 25. Response of the (8,7) nanosensor chirality to IL-6 in human serum.** Shift in emission center wavelength for (8,7) chirality of the IL-6 nanosensor in response to (A) 25 pg/mL IL-6. n=3, mean ± SD. Non-passivated (0.18 ± 0.02 nm), 50x BSA passivated (0.53 ± 0.13 nm), 50x PLK passivated (-0.68 ± 0.2 nm), and 50x NFDM passivated (0.14 ± 0.004 nm). For groups, Non-passivated and 50x BSA passivated nanosensor (0.34 nm, p=5E-2); Non-passivated and 50x PLK (0.87 nm, p=5.6E-4); Non-passivated and 50x NFDM (-0.04 nm, p=0.96). (B) Nanosensor response to 250 pg/mL IL-6. n=3, mean ± SD. Non-passivated (-0.2 ± 0.7 nm), 50x BSA passivated (0.25 ± 0.14 nm), 50x PLK passivated (-0.8 ± 0.15 nm), and 50x NFDM passivated (-0.15 ± 0.13 nm). For groups, Non-passivated and 50x BSA passivated nanosensor (0.45 nm, p=0.4); Non-passivated and 50x PLK (0.56 nm, p=0.25); Non-passivated and 50x NFDM (0.05 nm, p=1). (C) Nanosensor response to 2,500 pg/mL IL-6. n=3, mean ± SD. Non-passivated (0.32 ± 0.05 nm), 50x BSA passivated (0.41 ± 0.1 nm), 50x PLK passivated (-0.62 ± 0.53 nm), and 50x NFDM passivated (0.07 ± 0.05 nm). For groups, Non-passivated and 50x BSA passivated nanosensor (0.09 nm, p=1); Non-passivated and 50x PLK (0.94 nm, p=1.3E-2); Non-passivated and 50x NFDM (0.25 nm, p=0.58). (D) Nanosensor response to 25,000 pg/mL IL-6. n=3, mean ± SD. Non-passivated (0.2 ± 0.07 nm), 50x BSA passivated (0.09 ± 0.21 nm), 50x PLK passivated (-0.94 ± 0.24 nm), and 50x NFDM passivated (0.05 ± 0.12 nm). For groups, Non-passivated and 50x BSA passivated nanosensor (0.1 nm, p=0.87); Non-passivated and 50x PLK (1.1 nm, p=8.7E-4); Non-passivated and 50x NFDM (-0.14 nm, p=0.71).

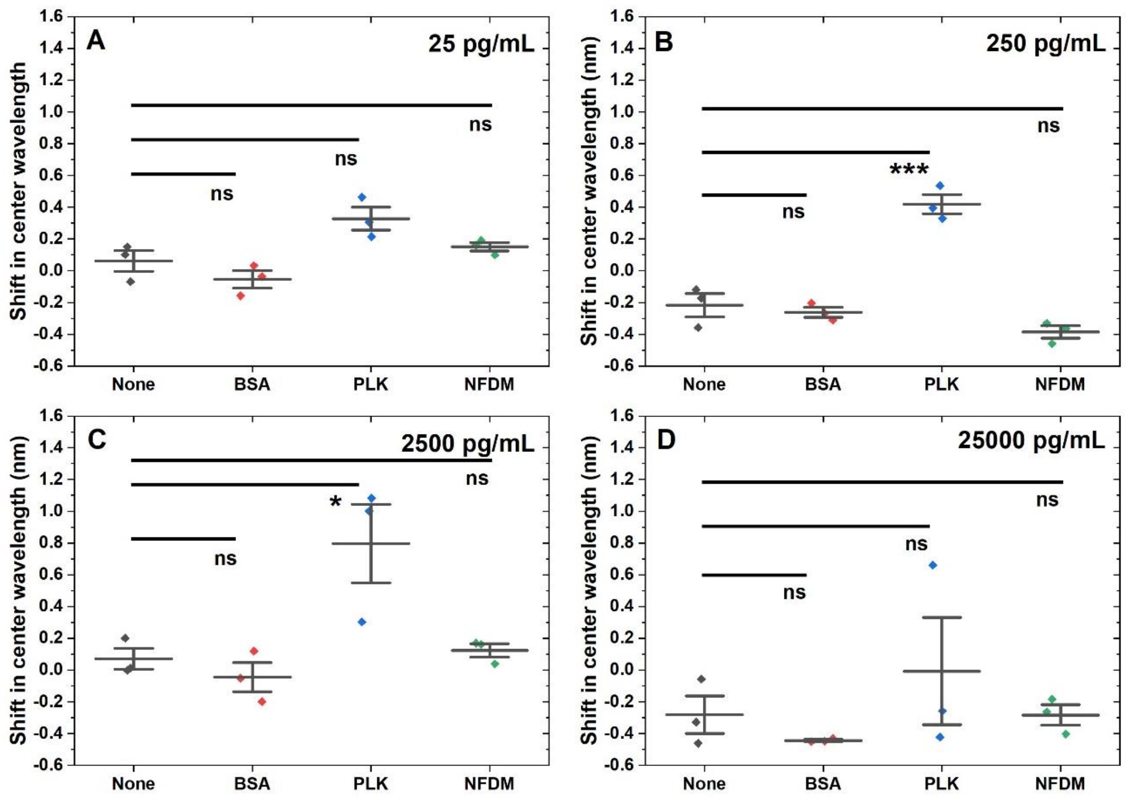


**Supplementary Figure 26. Response of the (7,6) nanosensor chirality to IL-6 in human serum.** Shift in emission center wavelength for (7,6) chirality of the IL-6 nanosensor in response to (A) 25 pg/mL IL-6. n=3, mean ± SD. Non-passivated (0.06 ± 0.12 nm), 50x BSA passivated (-0.05 ± 0.1 nm), 50x PLK passivated (0.33 ± 0.13 nm), and 50x NFDM passivated (0.15 ± 0.05 nm). For groups, Non-passivated and 50x BSA passivated nanosensor (0.12 nm, p=0.51); Non-passivated and 50x PLK (0.27 nm, p=0.07); Non-passivated and 50x NFDM (0.1 nm, p=0.68) (B) nanosensor response to 250 pg/mL IL-6. n=3, mean ± SD. Non-passivated (-0.2 ± 0.13 nm), 50x BSA passivated (-0.3 ± 0.1 nm), 50x PLK passivated (0.42 ± 0.11 nm), and 50x NFDM passivated (-0.4 ± 0.07 nm). For groups, Non-passivated and 50x BSA passivated nanosensor (0.04 nm, p=0.88); Non-passivated and 50x PLK (0.64 nm, p=3.04E-4); Non-passivated and 50x NFDM (0.17 nm, p=0.15) (C) nanosensor response to 2.5 ng/mL IL-6. n=3, mean ± SD. Non-passivated (0.07 ± 0.1 nm), 50x BSA passivated (-0.04 ± 0.16 nm), 50x PLK passivated (0.8 ± 0.43 nm), and 50x NFDM passivated (0.12 ± 0.07 nm). For groups, Non-passivated and 50x BSA passivated nanosensor (0.113 nm, p=0.9); Non-passivated and 50x PLK (0.73 nm, p=0.02); Non-passivated and 50x NFDM (0.05 nm, p=0.98) (D) nanosensor response to 25 ng/mL IL-6. n=3, mean ± SD. Non-passivated (-0.3 ± 0.1 nm), 50x BSA passivated (-0.44 ± 0.01 nm), 50x PLK passivated (-0.07 ± 0.6 nm), and 50x NFDM passivated (-0.3 ± 0.1 nm). For groups, Non-passivated and 50x BSA passivated nanosensor (0.16 nm, p=0.8); Non-passivated and 50x PLK (0.3 nm, p=0.49); Non-passivated and 50x NFDM (1E-3 nm, p=1)

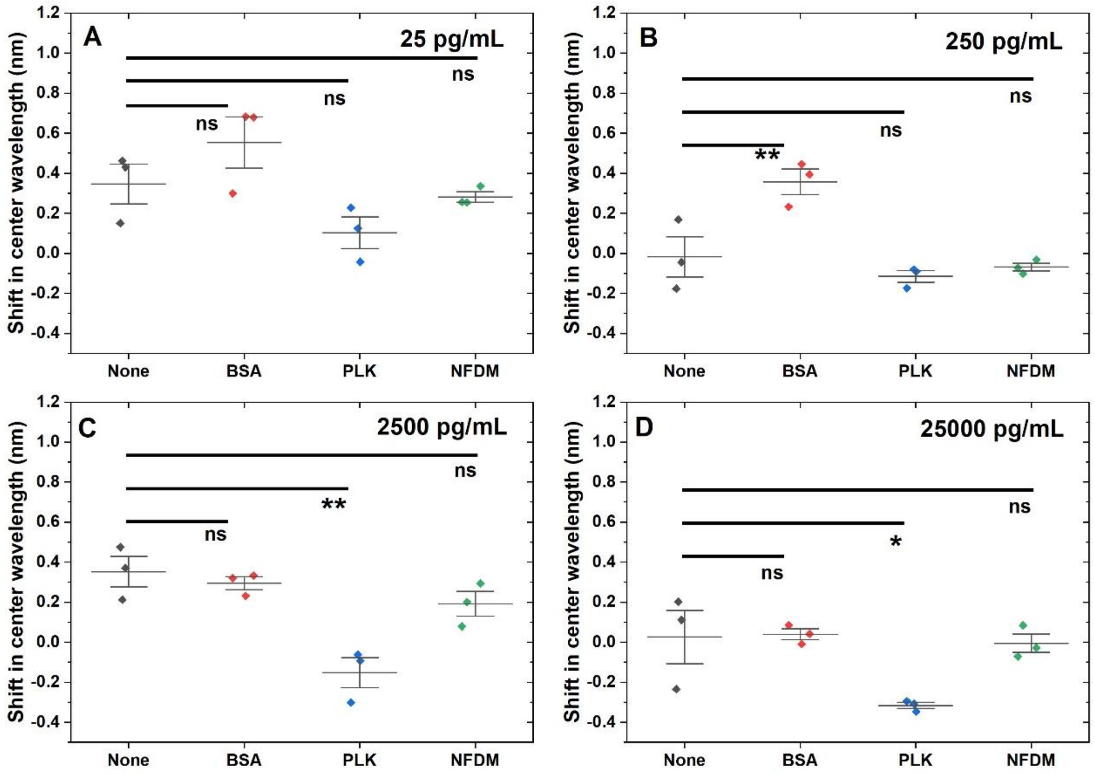


**Supplementary Figure 27. Response of the (9,4) nanosensor chirality to IL-6 in human serum.** Shift in emission center wavelength for (9,4) chirality of the IL-6 nanosensor in response to (A) 25 pg/mL IL-6. n=3, mean ± SD. Non-passivated (0.35 ± 0.02 nm), 50x BSA passivated (0.55 ± 0.22 nm), 50x PLK passivated (0.1 ± 0.14 nm), and 50x NFDM passivated (0.28 ± 0.02 nm). For groups, Non-passivated and 50x BSA passivated nanosensor (0.21 nm, p=0.4); Non-passivated and 50x PLK (0.24 nm, p=0.31); Non-passivated and 50x NFDM (0.1 nm, p=0.94). (B) Nanosensor response to 250 pg/mL IL-6. n=3, mean ± SD. Non-passivated (-0.02 ± 0.17 nm), 50x BSA passivated (0.36 ± 0.11 nm), 50x PLK passivated (-0.11 ± 0.05 nm), and 50x NFDM passivated (-0.07 ± 0.03 nm). For groups, Non-passivated and 50x BSA passivated nanosensor (0.37 nm, p=7.7E-3); Non-passivated and 50x PLK (0.1 nm, p=0.5); Non-passivated and 50x NFDM (0.05 nm, p=0.9). (C) Nanosensor response to 2,500 pg/mL IL-6. n=3, mean ± SD. Non-passivated (0.35 ± 0.13 nm), 50x BSA passivated (0.3 ± 0.06 nm), 50x PLK passivated (-0.15 ± 0.13 nm), and 50x NFDM passivated (0.2 ± 0.11 nm). For groups, Non-passivated and 50x BSA passivated nanosensor (0.06 nm, p=0.78); Non-passivated and 50x PLK (0.5 nm, p=1.1E-3); Non-passivated and 50x NFDM (0.16 nm, p=0.16). (D) Nanosensor response to 25,000 pg/mL IL-6. n=3, mean ± SD. Non-passivated (0.03 ± 0.05 nm), 50x BSA passivated (0.04 ± 0.05 nm), 50x PLK passivated (-0.32 ± 0.03 nm), and 50x NFDM passivated (-0.005 ± 0.11 nm). For groups, Non-passivated and 50x BSA passivated nanosensor (0.013 nm, p=1); Non-passivated and 50x PLK (0.34 nm, p=3.5E-2); Non-passivated and 50x NFDM (0.03 nm, p=1).

### Supplemental References


1. Iverson, N. M.; Barone, P. W.; Shandell, M.; Trudel, L. J.; Sen, S.; Sen, F.; Ivanov, V.; Atolia, E.; Farias, E.; McNicholas, T. P.; Reuel, N.; Parry, N. M. A.; Wogan, G. N.; Strano, M. S., In vivo biosensing via tissue-localizable near-infrared-fluorescent single-walled carbon nanotubes. Nature Nanotechnology **2013,** 8 (11), 873-880.
2. Babcock, J. J.; Brancaleon, L., Bovine serum albumin oligomers in the E- and B-forms at low protein concentration and ionic strength. Int J Biol Macromol **2013,** 53, 42-53.
3. Vincent, D.; Elkins, A.; Condina, M. R.; Ezernieks, V.; Rochfort, S., Quantitation and Identification of Intact Major Milk Proteins for High-Throughput LC-ESI-Q-TOF MS Analyses. PLoS One **2016,** 11 (10), e0163471.
